# Temporal Dynamics of Nucleus Accumbens Neurons in Male Mice During Reward Seeking

**DOI:** 10.1101/2024.09.27.615291

**Authors:** Terra A. Schall, King-Lun Li, Xiguang Qi, Brian T. Lee, William J. Wright, Erin E. Alpaugh, Rachel J. Zhao, Jianwei Liu, Qize Li, Bo Zeng, Lirong Wang, Yanhua H. Huang, Oliver M. Schlüter, Eric J. Nestler, Edward H. Nieh, Yan Dong

**Author notes:** Corresponding Author: Yan Dong, Address: Dept. Neuroscience, Univ. of Pittsburgh, A210 Langley Hall / 5^th^ & Ruskin Ave, Pittsburgh PA 15260. These authors contributed equally to this manuscript (Terra A. Schall, King-Lun Li, Xiguang Qi).

## Abstract

The nucleus accumbens (NAc) regulates reward-motivated behavior, but the temporal dynamics of NAc neurons that enable “free-willed” animals to obtain rewards remain elusive. Here, we recorded Ca^2+^ activity from individual NAc neurons when mice performed self-paced lever-presses for sucrose. NAc neurons exhibited three temporally-sequenced clusters, defined by times at which they exhibited increased Ca^2+^ activity: approximately 0, -2.5 or -5 sec relative to the lever-pressing. Dopamine D1 receptor (D1)-expressing neurons and D2-neurons formed the majority of the -5-sec versus -2.5-sec clusters, respectively, while both neuronal subtypes were represented in the 0-sec cluster. We found that pre-press activity patterns of D1- or D2-neurons could predict subsequent lever-presses. Inhibiting D1-neurons at -5 sec or D2-neurons at -2.5 sec, but not at other timepoints, reduced sucrose-motivated lever-pressing. We propose that the time-specific activity of D1- and D2-neurons mediate key temporal features of the NAc through which reward motivation initiates reward-seeking behavior.

## Introduction

Volitional reward-motivated behavior is not the mere act of taking a reward; it emerges through a sequence of preluding cognitive and other behavioral events. These events include processes like goal-engaging, action-initiating, distraction-blocking, and other cognitive/behavioral components, which operate collectively and sequentially to commit the animal towards making an operant response for a reward^1–3^. Neural underpinnings contributing to these pre-operant events remain largely unknown.

The nucleus accumbens (NAc) has long been recognized for its involvement in reward-motivated behaviors, specifically linking motivation to reward taking^4, 5^. Comprising >90% of the neuronal population in the NAc, principal medium spiny neurons (MSNs) are largely divided into two subpopulations, dopamine D1 receptor-expressing MSNs (D1-MSNs) and D2-MSNs, which are differentially involved in reward-motivated behavior^6–13^. Due to these cell-type-based differences and other heterogeneous features, NAc MSNs are thought to form distinct ensembles, each contributing to specific aspects of reward-motivated behavior^14–17^. Using in vivo recordings, rodent studies demonstrate that some NAc MSNs exhibit increased activity upon reward delivery and consumption, while some others preferentially respond to predictive cues preceding reward delivery^18, 19^, suggesting an ensemble-based organization of NAc MSNs across different temporal phases of motivational responses. Relevant to the pre-operant phase, elevated activities are observed in a population of NAc MSNs before a reward-conditioned task, while NAc lesions decrease the likelihood of rats to initiate operant responding for reward^20, 21^. These findings led us to examine an important yet unexplored question: whether NAc neurons exhibit unique activity patterns that contribute to the pre-operant cognitive and behavioral events in mice performing a well-learned, self-paced sucrose self-administration (SA) task.

Through GCaMP6m-mediated Ca^2+^ imaging, we observed that, during established sucrose SA, NAc neurons organized into three temporally distinct clusters, exhibiting activity increases at ∼0, ∼-2.5 or ∼-5 sec relative to the time of the lever-press. We show that the -5- and -2.5-sec clusters comprised preferentially D1-versus D2-neurons, respectively, while both D1- and D2-neurons constituted the 0-sec cluster. In addition, the activity patterns of D1-neurons at ∼-5 sec and D2-neurons at ∼-2.5 sec provided heightened prediction accuracy for subsequent lever-presses. Optogenetic inhibition of D1-neurons at -5 sec or D2-neurons at -2.5 sec, but not at other timepoints, reduced subsequent lever-pressing for sucrose. Together, these findings offer insight into the sequential activity changes of NAc D1- and D2-neurons that link reward motivation to behavioral output.

## Results

### Three clusters of NAc neurons

To detect the temporal dynamics of individual NAc neurons in freely-moving mice, we stereotaxically injected GCaMP6m-expressing AAV9 into the NAc and installed a GRIN lens above the injection site (Fig. 1AB). While the medial shell of the NAc was targeted, portions of the NAc core were also likely included in the sampling and viral manipulations due to the small size of the mouse NAc (Fig. S1AB). Six weeks later, we trained these mice with an overnight (12-h; see Methods) sucrose SA session, followed by an 11-d SA procedure (1 h/session/d), during which the mice were allowed to move freely and lever-press for a sucrose solution (10%) (Fig. 1CD). During the first 20 min of selected SA sessions, we recorded GCaMP6m-mediated Ca^2+^ signals continuously through a miniaturized fluorescence microscope (Miniscope)^22^ and extracted Ca^2+^ transients from individual NAc neurons (Fig. 1E-G).

**Figure 1.**
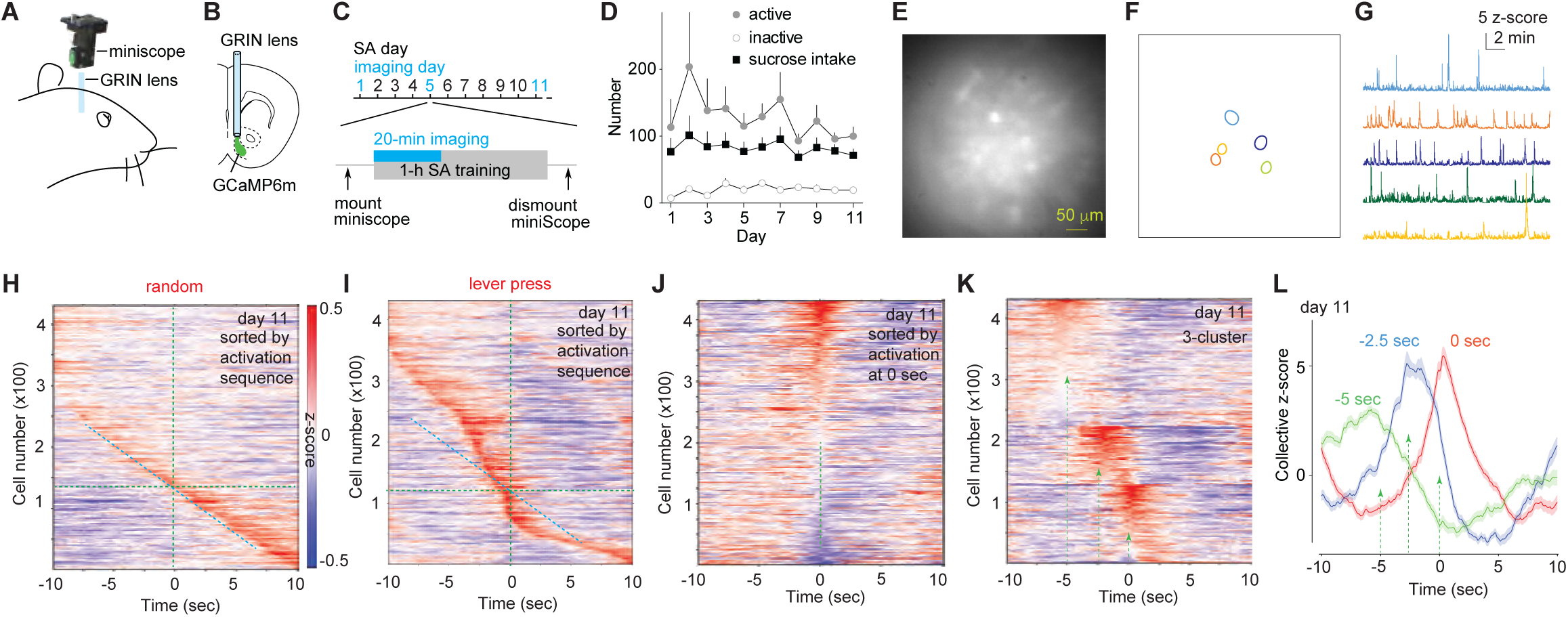
Three neuronal clusters in sucrose-motivated operant responding in wildtype mice. **A,B** Diagrams showing installation of Miniscope and GRIN lens to image activities of NAc neurons in vivo. **C** Schematic showing the 11-d sucrose SA procedure and imaging periods during training sessions. **D** Summaries of sucrose intake and operant responses during SA in the 7 mice used for in vivo imaging. Data point presenting mean ± standard error. **E** Example in vivo image of GCaMP signals in NAc neurons. **F,G** Example neurons selected from **E** (**F**) for extraction of Ca^2+^ transients (**G**). **H,I** Sequence-based sorting of neurons with increased activities during randomly selected 20-sec time windows (control, **H**) or during time windows with operant responding for sucrose on d 11 (**I**). Vertical dashed lines indicate the 0-sec timepoint. Horizontal and oblique lines (with same slope) are drawn to aid visualization. **J** Timing-based sorting of neurons with increased activities to 0 sec showing the 0-sec cluster as well as some neurons with increased activities during the pre-press phase. **K** K-means clustering detected three clusters that best described the temporal dynamics of NAc neurons during the 20-sec lever-press time window. Dashed lines/arrows indicate -5, -2.5 and 0 sec. **L** Summary of the collective z-scores of activated neurons over the 20-sec lever-press time window showing three temporally distinct neuronal clusters. Dashed lines/arrows indicate -5, -2.5 and 0 sec. Source data are provided in the Source Data file associated with this manuscript. Lines and shades presenting means and standard errors, respectively.

To quantify activity changes of individual neurons, we computed z-scores of Ca^2+^ traces (see Methods). We defined the time of lever-press as the 0-sec timepoint and created a dataset that extracted z-scores of individual neurons between 10 sec before and 10 sec after each lever-press from 7 wildtype mice (Fig. S1B). We first performed sequence-based sorting, which aligned neurons by the time they exhibited initial (Fig. 1HI) or peak (Fig. S1CD) activity increases. In a control dataset, in which neuronal activities were sampled during a randomly selected 20-sec time window (trial) without lever-press, neurons were distributed evenly over time, indicating a quasi-random activity pattern of NAc neurons when mice moved freely without involving motivated sucrose seeking (Figs. 1H;S1C). During lever-press trials on SA d 11, more neurons were detected before or around the lever-press timepoint (Figs. 1I;S1D), suggesting that select populations of NAc neurons synchronized their activities during the pre- and on-press phases.

To explore the timing of synchronized neuronal activities during the 20-sec trials, we performed a timing-based sorting, which aligned neurons to a specific timepoint based on their increases in z-scores. When sorting neurons with increased activities to 0 sec, we observed a 0-sec neuron cluster as well as a substantial number of neurons with increased activities before 0 sec (Fig. 1J), suggesting the existence of pre-press neuron clusters. We thus performed K-means clustering to identify the timepoints at which NAc neurons would exhibit synchronized activity increases. This analysis revealed three neuron clusters, exhibiting increased activities at ∼-5, ∼-2.5, and ∼0 sec, respectively (Fig. 1K). We also examined clustering neurons into a smaller or larger number of clusters, but found that three clusters performed the best (Fig. S1E-G,J).

Further analysis revealed that, within each of these three clusters, some neurons exhibited increased activities with their mean z-scores >0 over a 1-sec range around the cluster center timepoint, while others displayed mean z-scores <0. Based on these activity properties, we operationally defined them as “activated” versus “nonactive” neurons, respectively (Fig. S1H). Thus, activated neurons may represent a neuronal population, or a functional ensemble, that is selectively involved in constructing the pre-press temporal dynamics. To illustrate the time-dependent activity changes of the three clusters, we plotted collective z-scores (integration of number and activity intensity of neurons) of either activated neurons (Fig. 1L) or top 50% of neurons ranked by activity (see Methods; Fig. S1I), with both showing similar temporal dynamics for each of the three clusters. Together, these clusters may represent three at least partly distinct functional ensembles that sequentially contribute to the cognitive-behavioral events during the pre- or on-press phases of sucrose-motivated behavior.

### D1- versus D2-neurons

Representing the two major subtypes of principal neurons in the NAc, D1- and D2-MSNs may contribute differentially to the three clusters related to sucrose SA. To explore this, we injected AAV-Flex-GCaMP6m in the NAc of D1-Cre or D2-Cre mice, which resulted in GCaMP6m expression in D1- or D2-neurons, respectively. Approximately 6 weeks later, we monitored activities of individual neurons in these mice during the 11-d sucrose SA procedure (Fig. S2A-D).

On SA d 11, sequence-based sorting revealed prominent portions of D1-neurons exhibiting increased activities during the pre- and on-press phases (Fig. 2A). Timing-based sorting of D1-neurons to 0 sec showed a 0-sec cluster, as well as a substantial percentage of neurons exhibiting increased activities during the pre-press phase (Fig. 2B). Similarly, populations of D2-neurons exhibited activity increases during both the pre- and on-press phases on SA d 11, (Fig. 2CD).

**Figure 2.**
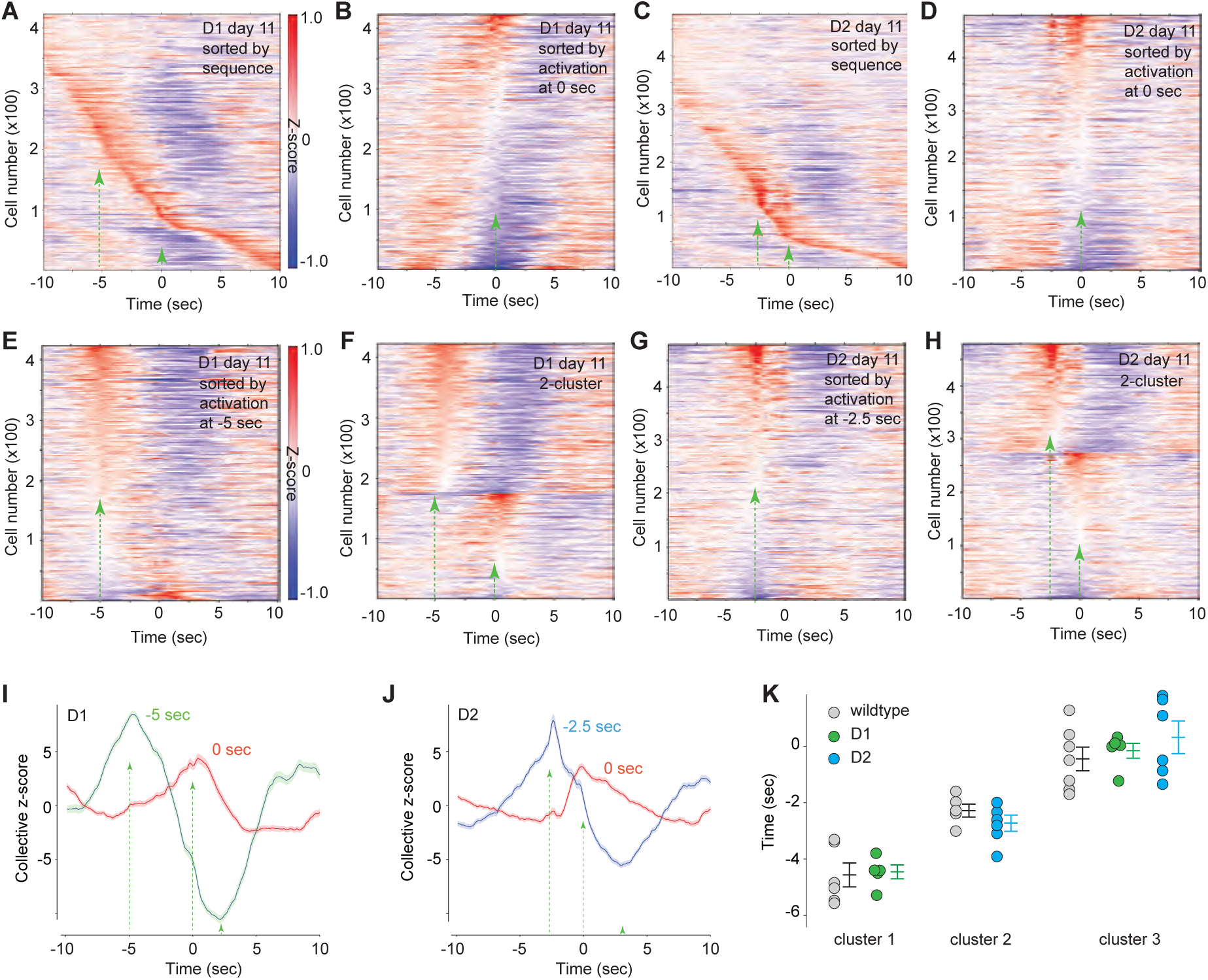
Differential contributions of D1- and D2-neurons to NAc clusters. **A** Sequence-based sorting of D1-neurons with increased activities on sucrose SA d 11. Dashed lines/arrows indicate timepoints of 0, -2.5 or -5 sec in this and following panels. **B** Timing-based sorting of D1-neurons with increased activities to 0 sec. **C** Sequence-based sorting of D2-neurons with increased activities on SA d 11. **D** Timing-based sorting of D2-neurons with increased activities to 0 sec. **E** Timing-based sorting of D1-neurons with increased activities to -5 sec. **F** K-means clustering results showing two clusters of D1-neurons. **G** Timing-based sorting of D2-neurons with increased activities to -2.5 sec. **H** K-means clustering showing two clusters of D2-neurons. **I,J** Collective z-scores of activated D1-neurons (**I**) or D2-neurons (**J**) in each cluster showing the temporal dynamics over the 20-sec time window. Lines and shades presenting means and standard errors, respectively. **K** Summaries showing the mean timepoints of K-means clustering-defined clusters of D1-neurons (in sec: -4.7 ± 0.3; -0.2 ± 0.2; n = 5) or D2-neurons (in sec: -2.8 ± 0.3; 0.3 ± 0.2; n = 6), as well as neurons from wildtype mice (in sec: -4.6 ± 0.4; -2.3 ± 0.2; -0.4 ± 0.4; n = 7). Middle and outer lines presenting means and standard errors, respectively. Source data are provided in the Source Data file associated with this manuscript.

Over the 20-sec trial, we found more D1- and D2-neurons that exhibited high activity (i.e., >95% of the mean z-scores of all neurons) at ∼-5 and -2.5 sec, respectively, compared to other timepoints (Fig. S2E-H). Furthermore, rather than the three neuronal clusters deduced in wildtype mice, the cluster-wise assessment of cluster stability revealed that two clusters best captured the organization of either D1-neuron or D2-neuron activities during the pre- and on-press phases (Fig. S2IJ). Based on the timing of the two pre-press clusters described in wildtype mice, we performed timing-based sorting to align D1- and D2-neurons with increased activities to either -2.5 or -5 sec. We found that -5 sec better aligned the pre-press activities of D1-neurons (Fig. 2E; S2KL). K-means clustering likewise revealed that the activities of D1-neurons were described by two clusters centered at ∼-5 and ∼0 sec (Fig. 2F). On the other hand, timing-based sorting to -2.5 sec better aligned the pre-press activities of D2-neurons compared to -5 sec (Fig. 2G; S2MN), with K-means clustering also identifying two clusters of D2-neurons centered at ∼-2.5 and ∼0 sec (Fig. 2H).

As observed for neurons in wildtype mice, D1- or D2-neurons were both composed of activated and nonactive portions in each cluster (Fig. S2OP). Time-dependent changes in collective z-scores of activated neurons revealed distinct temporal dynamics of the D1-neuron versus D2-neuron clusters during the pre-press phase (Fig. 2IJ). As a further validation, we pooled all activated D1- and D2-neurons together for K-means clustering analysis. This reconstituted populations of both neuronal subtypes once again revealed three clusters with the temporal dynamics consistent to those seen in wildtype mice (Fig. S2Q). Importantly, the -5- and -2.5-sec clusters grouped by K-means clustering from pooled neurons were enriched in D1- and D2-neurons, respectively (Fig. S2R).

The three timepoints (∼-5, ∼-2.5, and ∼0 sec) above were estimated based on neurons pooled from all mice in each group. K-means clustering of individual mice revealed between-subject variations, but the deductions remained consistent with the mean timepoints (Fig. 2K).

Taken together, our findings show that D1-neurons and D2-neurons differentially contribute to the two sequential clusters during the pre-press phase, which may contribute to a sequence of cognitive-behavioral events that leads to operant responses for sucrose. Meanwhile, both D1- and D2-neurons contribute to the 0-sec cluster, corresponding to the lever-press action and sucrose consumption.

### Populational activity patterns

To test whether the pre-press neuronal activity can predict lever-pressing, we performed a variant of principal component analysis (PCA)^23^, in which we built our PCA matrix on a trial-averaged 30-sec time window, centered around lever-presses (Fig. S3A). The first two principal components (PCs) captured >30% of the variance of trial-averaged data from D1- or D2-neurons, as well as neurons in wildtype mice (Fig. S3B).

For each mouse on SA d 11, we plotted the PC1 and PC2 values over the 20-sec trials with lever-pressing, using the PC plot of randomly selected 20-sec trials as baseline controls. In an example D1-Cre mouse, the lever-press and baseline control PC data points appeared largely overlapping at -10 sec but separated at 0 sec (Figs. 3AB; S3CD). To quantify this separation, we identified the centroids of lever-press versus non-lever-press data points in each mouse, measured the Euclidian distance between the 2 centroids, and normalized it to the centroid distance between two baseline controls in the same mouse to obtain the relative distance (Figs. 3AB;S3CD). In D1-neurons, the relative distance of lever-press exhibited a peak at ∼-5 sec, followed by larger increases ∼0 sec (Figs. 3C). In D2-neurons, the relative distance of lever-press exhibited an increase at ∼-2.5 sec, followed by larger increases at ∼0 sec (Figs. 3D). When assessed by the bootstrapped confidence interval (BCI; see Methods) analysis, although most intervals (n = 100 samples x 5-fold points = 500 / interval) of active lever-press trials exhibited higher relative distance values compared to control trials, the magnitudes of confidence around -5 sec (t value interval, 22. 5 – 27.2) were not overlapping with and were significantly higher than -2.5 sec (7.6 – 11.9) in D1-neurons, as were D2-neurons at -2.5 sec (32.7 – 38.1) compared to -5 sec (5.6 – 9.7). When repeating this in a higher dimensional space with 10 PCs, we observed similar results (Fig. S3EF). These results confirm that D1- and D2-neurons display distinct temporal dynamics during the pre-press phase.

**Figure 3.**
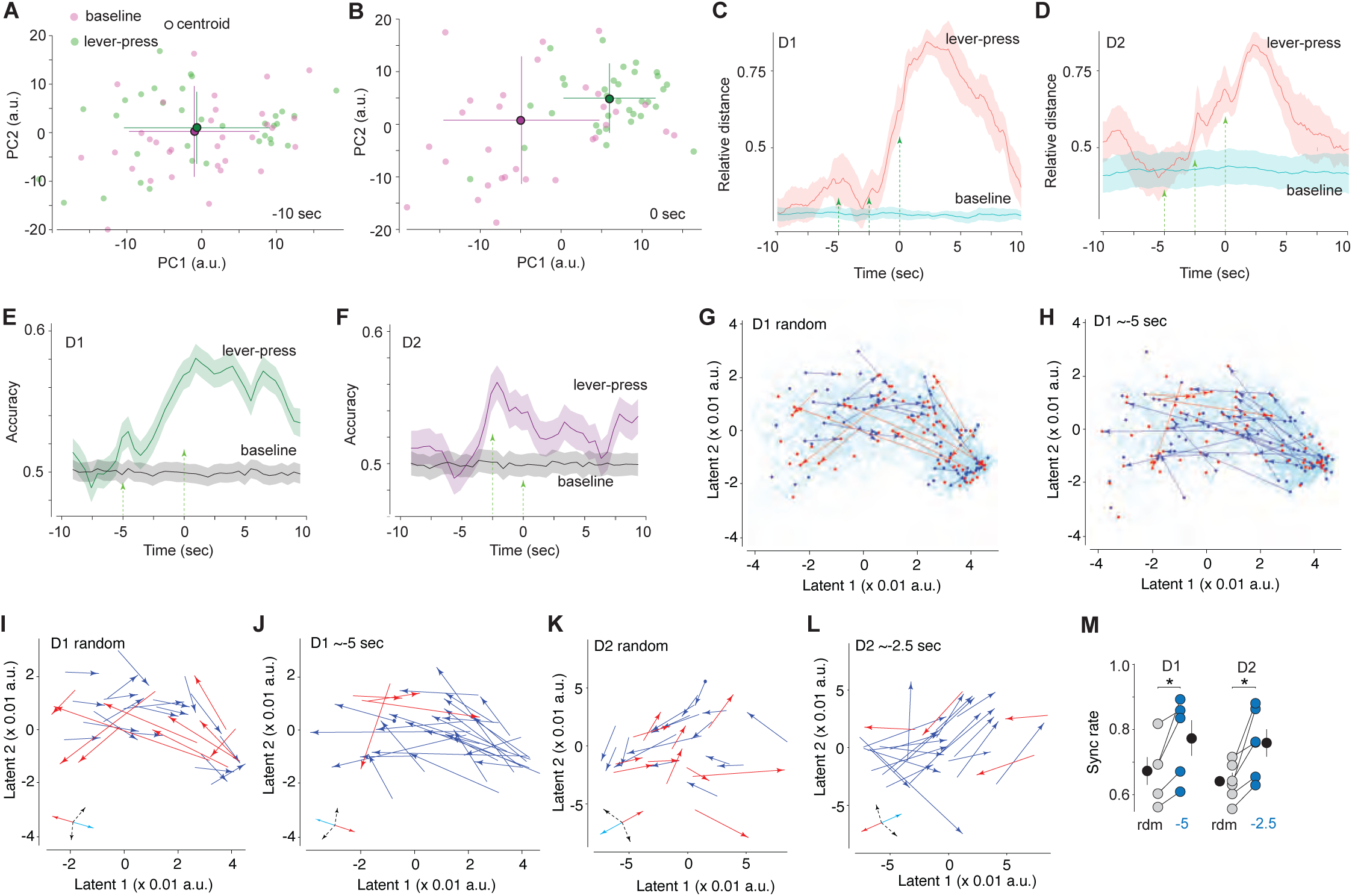
Populational activities of D1- and D2-neurons before lever-pressing for sucrose. **A,B** Two-dimensional PCA projections of NAc neuronal activities at -10 (**A**) and 0 sec (**B**) from an example mouse. Each dot represents neuronal activities of a single trial (purple, baseline; green, lever-press; dark dots, centroids; crosslines, standard deviation). Center dots and lines representing means and standard deviations, respectively. **C,D** The distance of PCA centroids of D1-neurons (**C**) or D2-neurons (**D**) between lever-press and baseline trials. Solid lines, mean distance of all tested mice; shade, standard errors. Dashed lines/arrows indicate timepoints of 0, -2.5 or -5 sec. **E,F** Support Vector Machine accuracy of lever-press prediction based on two latents extracted from D1-neuron (**E**) and D2-neuron (**F**) activities using MIND. Solid lines, mean accuracy of all tested mice, shade, standard errors. **G,H** Example mouse (D1-Cre) showing two-latent projections of D1-neuron activity of all lever-press trials. Each dot indicates the two-latent position of neurons at a single timepoint in a trial. Solid lines connecting two sets of latent coordinates of neurons at the times of (t – 1.5 sec) and (t) in each trial. “t” was either randomly selected (**G**) or set to ∼-5 sec (**H**), with arrows indicating the direction from (t – 1.5 sec) to (t), to represent the trajectories through which neurons traveled over the 1.5-sec period. Arrows at angles within ±75° with the mean direction of all arrows were coded in blue and others were coded in red. **I,J** Arrow plots of neural trajectories of D1-neurons over the 1.5-sec periods with randomly selected t (**I**) or t = ∼-5 sec (**J**) from an example D1-Cre mouse. **K,L** Arrow plots of neural trajectories of D2-neurons over the 1.5-sec periods with randomly selected t (**K**) or t = ∼-2.5 sec (**L**) from an example D2-Cre mouse. *Insets* in **I-L**: blue arrow, mean direction calculated as the mean angle of all arrows; red arrow, opposite direction of the blue arrow; black arrows, describing the range of included angle of ±75° with the mean direction. **M** Summaries showing synchronized trajectories of D1-neurons to ∼-5 sec (p = 0.03 rdm versus -5, two-sided paired t-test) and D2-neurons to ∼-2.5 sec (p = 0.01 rdm versus -2.5, two-sided paired t-test). Each dot representing data from a single mouse (5 D1-cre and 6 D2-cre mice). Black dots and vertical lines representing means and standard errors, respectively. *p < 0.05. Source data are provided in the Source Data file associated with this manuscript.

The linear assumption of PCA may omit some nonlinear aspects of neuronal dynamics, so we used MIND^24^ to conduct nonlinear dimensionality reduction. Using the 7-dimensional embedding of the manifold (Fig. S3G), we took the first two dimensions and built a series of Support Vector Machine-based classifiers^25, 26^ to test whether neuronal activities during the pre-press phase predict the action of lever-pressing. Our results show that the ability of Support Vector Machine classifiers to predict the upcoming lever-press increased at ∼-5 sec for D1-neurons [t-value BCI interval (n = 100 samples/intervals), 7.3 – 12.2] and at ∼-2.5 sec for D2-neurons [t-value BCI interval (n = 100 samples/intervals), 16.0 – 21.2], respectively (Fig. 3EF).

To further characterize the conserved neural dynamics of D1- and D2-neurons that may predict future lever-presses, we examined the flow of neural trajectories. In two-latent dimensions from MIND outputs, we plotted the coordinates of D1- or D2-neurons at individual timepoints over the 20-sec trials, and used arrows that connected the two timepoints (1.5 sec apart) to describe the neural trajectories through which neurons traveled (Figs. 3G-L;S3H-J). Our results show that the trajectories of D1-neurons to ∼-5 sec, and the trajectories of D2-neurons to either -5 or -2.5 sec, converged better than the trajectories to randomly selected control timepoints (Figs. 3M; S3J). These results confirm – with some caveats (see Discussion) – the organized temporal dynamics of D1- and D2-neurons during the pre-press phase.

### Behavioral correlates

To explore whether the two pre-press clusters are correlated with mice’s movements, we used DeepLabCut (DLC)^27^ to trace and analyze the motion of different body parts of the mouse during the 20-sec lever-press trials, using trials without lever-press as baseline controls (Fig. 4AB). Among the three major body parts (head, main body, and tail), the movement velocity of heads (averaged from 7 D1-Cre and 7 D2-Cre mice) exhibited relatively clear changes during pre-press phases, with an initial increase at ∼-5 sec and peak at ∼2.5 sec (Fig. S3K-M). Furthermore, over the 20-sec lever-press trials, the activity intensities (z-scores) of D1-neurons and D2-neurons were correlated positively with the velocities of head movement, and these correlations were more prominent for D1-neurons at -5 sec and D2-neurons at -2.5 sec, compared to other timepoints (Fig. S3N-Q). These correlations suggest a potential role of NAc clusters in either locomotion or cognitive states that are behaviorally manifested by head movements. Nonetheless, within the operant chamber, locations (assessed by head locations) of either D1-Cre or D2-Cre mice were rather random at -7.5 and -5 sec, but moved close to the final responding zone (i.e., mouse’s location at -0.5 sec) at -2.5 sec (Fig. S3R-X). Moreover, the synchronized movement trajectories toward the final responding zone were not detected at -5 sec, but at -2.5 sec (Fig. S3Y-CC). These results suggest that the increase in head movement at -5 sec may not signal locomotion, but reflect certain cognitive states.

**Figure 4.**
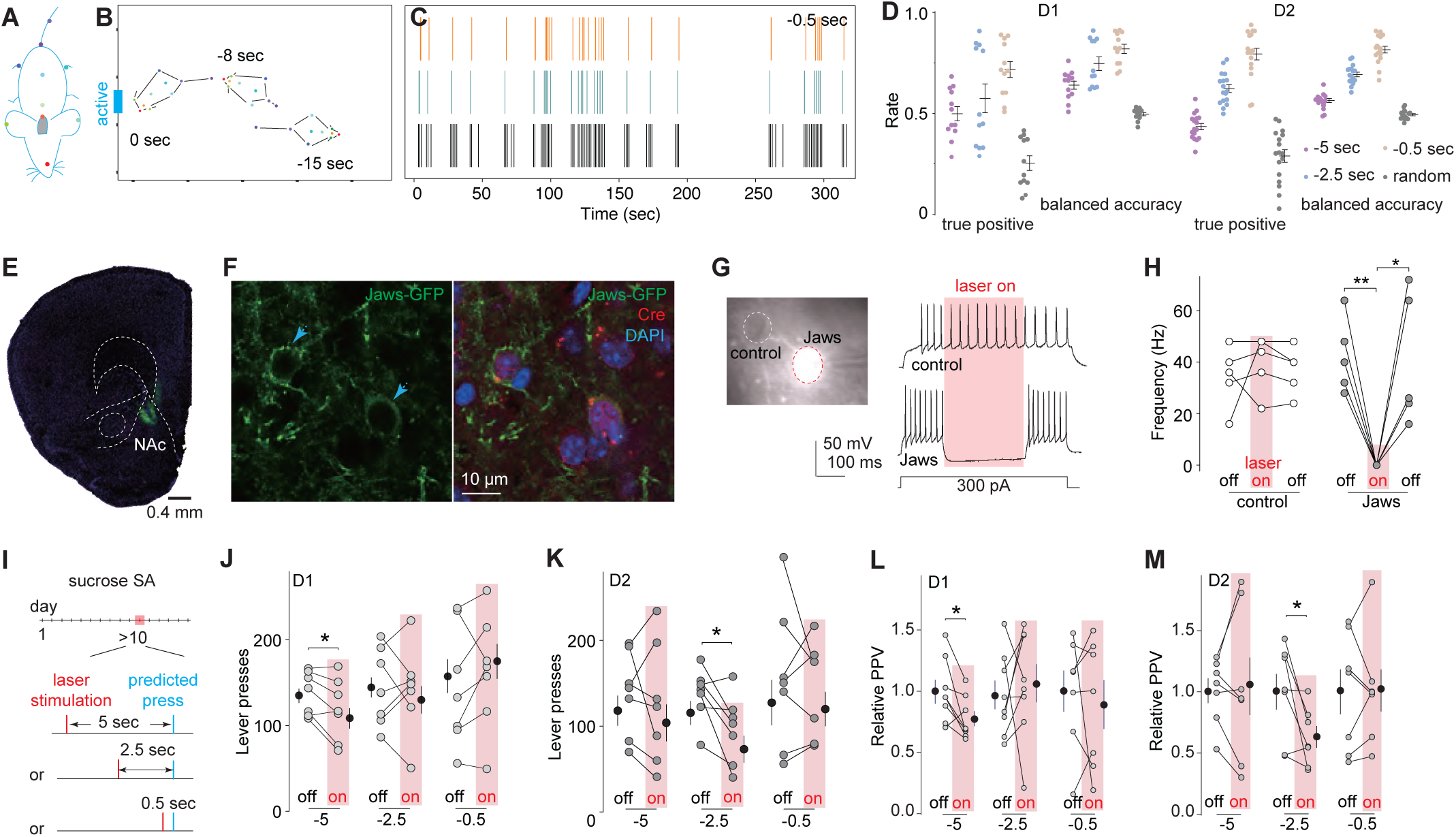
Interfering with the pre-press temporal dynamics of NAc neurons reduces sucrose-motivated operant responses. **A** Illustration showing positions where tracing dots were assigned on mouse’s body in DeepLabCut (DLC) analysis. **B** Example DLC tracing of a mouse before and upon lever-pressing for sucrose. **C** Example DLC prediction of lever-presses for sucrose. Upper, yellow bars show the times when actual lever-presses occurred. Middle, blue bars show the times that were 0.5 sec before yellow bars, the timepoints where the prediction should be made. Lower, black bars show DLC-predicted timepoints that were 0.5 sec before the potential lever-presses. **D** Summaries showing the true positive rate and balanced accuracy of the 5-sec, -2.5-sec and -0.5-sec DLC models, with the control of random prediction (D1 true positive: - 5 sec 0.50 ± 0.04, - 2.5 sec 0.57 ± 0.07, -0.5 sec 0.71 ± 0.04, random 0.25 ± 0.04, n = 12; D1 balanced accuracy: -5 sec 0.64 ± 0.02, - 2.5 sec 0.75 ± 0.03, -0.5 sec 0.82 ± 0.02, random, 0.50 ± 0.01, n = 12; D2 true positive: -5 sec 0.44 ± 0.02, - 2.5 sec 0.63 ± 0.02, -0.5 sec 0.80 ± 0.03, random 0.28 ± 0.04, n = 17; D2 balanced accuracy: -5 sec 0.57 ± 0.01, - 2.5 sec 0.70 ± 0.01, -0.5 sec 0.82 ± 0.02, random, 0.50 ± 0.01, n = 17). Each dot (n) indicates a session of a mouse. Middle and outer lines presenting means and standard errors, respectively. **E** Example slice showing intra-NAc viral-mediated expression of Jaws. **F** Images from an example D1-Cre mouse showing that Cre-positive neurons co-expressed Jaws. Note that Jaws was preferentially expressed on the plasma membrane. **G** Example neurons with or without Jaws expression in the NAc slice (left) and example recordings (right) showing that laser activation of Jaws silenced action potential firing in a Jaws-expressing neuron, but not control neuron. **H** Summaries showing that laser activation of Jaws decreased the frequency of evoked action potential firing in NAc neurons that expressed Jaws without affecting control neurons (in Hz; control: off_1_ 34.4 ± 5.3, on 39.6 ± 4.9, off_2_ 36.8 ± 4.1, n = 5 neurons from 3 mice, F_2,12_ = 0.43, p = 0.55; Jaws: off_1_ 42.0 ± 5.2, on, 0.0 ± 0.0, off_2_, 37.7 ± 9.7, n = 6 neurons from 3 mice, F_2,15_ = 19.06, p = 0.00; p = 0.00 off_1_ vs on, p = 0.04 off_2_ vs on; one-way ANOVA repeated measures followed by Bonferroni’s posttest). **I** Schematic showing optogenetic inhibition of NAc neurons at -5, -2.5 or -0.5 sec before the predicted lever-press after 10 d of sucrose SA. **J,K** Summaries showing that optogenetic inhibition of NAc D1-neurons at -5 sec (-5 sec: off 138.0 ± 9.6, on 119.3 ± 13.7, t_6_ = 2.7, p = 0.03; -2.5 sec: off 144.8 ± 17.1, on 142.4 ± 19.3, t_6_ = 0.2, p = 0.88; -0.5 sec: off 146.2 ± 26.5, on 163.0 ± 25.1, t_6_ = 1.09; p = 0.32, n = 7 mice; paired t-test; **J**) or D2-neurons at -2.5 sec (-5 sec: off 137.8 ± 18.6, on 124.4 ± 26.8, t_6_ = 0.67, p = 0.53; -2.5 sec: off 136.8 ± 11.7, on 96.1 ± 15.1, t_6_ = 2.86, p = 0.03; - 0.5 sec: off 155.4 ± 31.9, on 145.0 ± 21.1, t_6_ = 0.45, p = 0.67; n = 7 mice; two-sided paired t-test; **K**), but not other timepoints, decreased the number of lever-presses for sucrose. **L,M** Summaries showing decreased relative Positive Predictive Value (PPV) of DLC-based prediction upon optogenetic inhibition of NAc D1-neurons at -5 sec (-5 sec, off 1.00 ± 0.09, on 0.76 ± 0.06, t_7_ = 2.8, n = 8, p = 0.03; -2.5 sec, off 1.00 ± 0.12, on 1.09 ± 0.16, t_7_ = 0.4, n = 8 mice, p = 0.67; -0.5 sec, off 1.00 ± 0.17, on 0.90 ± 0.20, t_6_ = 0.5, n = 7 mice, p = 0.65; paired t-test; **L**) or D2-neurons at -2.5 sec (5 sec, off 1.00 ± 0.10, on 1.03 ± 0.23, t_6_ = 0.2, n = 7, p = 0.82; -2.5 sec, off 1.00 ± 0.15, on 0.63 ± 0.09, t_6_ = 2.7, n = 7, p = 0.04; -0.5 sec: off 1.01 ± 0.17, on 1.01 ± 0.17, t_6_ = 0.13, n = 7, p = 0.93; two-sided paired t-test; **M**), but not other timepoints. *p < 0.05; **p < 0.01. Source data are provided in the Source Data file associated with this manuscript.

To directly examine the roles of specifically-timed NAc clusters in launching the final act of lever-pressing for sucrose, we optimized a DLC-based algorithm to predict individual lever-presses during SA sessions. Our plan was to use in vivo optogenetics to interfere with the activity of NAc neurons in vivo at -5, -2.5 or -0.5 sec, and examine whether these cluster-targeted interferences influence the likelihood of subsequent lever-presses. The first two timepoints (-5 and -2.5 sec) correspond to the two pre-press neuronal clusters. The -0.5-sec timepoint corresponds to the rising phase of the 0-sec cluster (Figs. 1L;2IJ). We did not target 0 sec per se, at which point the action of lever-pressing was already under way.

We trained the DeepLabCut algorithm using the combined information of mouse’s postures, positions, and movements during the pre-press phase (Figs. 4AB;S4A). For each intended timepoint, the prediction was made 1 sec earlier, using the 5-sec video immediately before the prediction-making timepoint, such that there was sufficient lead time for the computer to process positive predictions and trigger optogenetic inhibitions exactly at the intended timepoint. For all three timepoints for the following experiments in both D1- and D2-Cre mice, the prediction efficacy was reasonably high, with both the true positive rate and balanced accuracy (see Methods) >50%, substantially higher than randomized predictions (Figs. 4CD, S4BC). Thus, ∼50% of the actual lever-presses could be captured by our DeepLabCut algorithm for the -5-sec timepoint. The prediction efficacy increased further for the -2.5-sec and -0.5-sec timepoints. Taken together, our DeepLabCut algorithm predicts a portion of, but not all, lever-presses.

To interfere with neuronal clusters in vivo, we expressed Jaws, a chloride pump for optogenetic inhibition upon 635-nm laser stimulation^28^, selectively in NAc D1- or D2-neurons by bilateral injection of AAV8-Flex-Jaws in D1-Cre or D2-Cre mice (Fig. 4EF;S4D). To target a particular cluster with minimal impact on other clusters, we verified a short inhibition duration, 250 ms, which was expected to interfere with, but not abolish, activities of the intended cluster. To verify the inhibition efficacy of this Jaws approach, we made whole-cell current-clamp slice recordings from NAc neurons with or without Jaws expression. A 500-ms depolarizing current step evoked persistent firing of action potentials in both Jaws-expressing (visualized by co-expressed GFP) and control (without expressing Jaws) neurons, while a laser stimulation (250 ms) selectively suppressed the evoked action potential firing in Jaws-expressing neurons (Fig. 4GH).

Through pre-installed optical fibers (Fig. S4MN), we applied this short-duration optogenetic inhibition to Jaws-expressing NAc neurons at the -5-sec, -2.5-sec or -0.5-sec timepoint during a 1-h sucrose SA session after >10 d of sucrose SA training (Fig. 4I). Compared to sessions without in vivo optogenetic stimulation (the “off” session), the same D1-Cre mice during the sessions with laser inhibition (the “on” session; 5-10 mW x 250 ms, bilateral) of NAc D1-neurons at -5 sec, but not at -2.5 or -0.5 sec, exhibited reduced levels of subsequent lever-presses for sucrose, as well as operant responses to the inactive lever without changes in responding accuracy [active/(active + inactive)] (Fig. 4J;S4E-G). Analogous results were obtained upon laser inhibition of NAc D2-neurons but with peak effects at -2.5 sec and not at -5 or -0.5 sec (Fig. 4K, S4I-K). In an additional analysis, we examined the lever-presses that were not predicted by DLC, thus without time-contingent optogenetic stimulation during the “on” session. In both D1-Cre and D2-Cre mice, the numbers of unpredicted lever-presses were similar between the “off” and “on” sessions (Fig. S4HL). As such, the basic sucrose-motivated operant responding was similar across sessions, but reduced by specifically-timed inhibition of D1- or D2-neurons.

The accuracy of DeepLabCut predictions can be quantified by the Positive Predictive Value [PPV; = true positives/(true positives + false positives)]. In these mice, the predictions were made before laser inhibition. As such, laser inhibition did not affect the number of predicted lever-presses but should decrease the PPV if the inhibition reduced the number of actual lever-presses. Indeed, the relative PPV (normalized to sessions without laser inhibition) was decreased upon optogenetic inhibition of NAc D1-neurons at -5 sec (Fig. 4L) or D2-neurons at -2.5 sec (Fig. 4M), but not any other timepoints for either of these neuronal types.

Taken together, these results establish that the -5-sec D1-neuron and -2.5-sec D2-neuron clustering activities represent two distinct steps in the sequence of neuronal events by which the NAc contributes to the initiation of sucrose-motivated operant responses.

## Discussion

In well-learned reward-motivated behaviors, the act of reward taking can be regarded as the final behavioral output of a sequence of cognitive-behavioral events. We propose that the clusters of neuronal dynamics in the NAc revealed by the present study contribute importantly to the circuit basis for some of these preluding events that launch reward seeking and taking behavior.

### The three clusters

Constituting >90% of neuronal populations in the NAc, MSNs were expected to be predominantly represented in the NAc clusters in our current study. Without intrinsic pace-making mechanisms, activation of NAc MSNs is driven primarily by glutamatergic inputs^29^. A single NAc MSN receives convergent inputs from the hippocampus, amygdala, several subregions of prefrontal cortex, and other limbic/paralimbic regions that encode different aspects of reward incentive^30, 31^. These inputs often adjacently synapse on the same MSN dendrites, together with heterosynaptically aligned dopaminergic and other monoaminergic presynaptic terminals^30, 32, 33^. Furthermore, these glutamatergic and monoaminergic inputs influence NAc interneurons, such as fast-spiking and cholinergic interneurons, which, in turn, regulate the firing pattern of MSNs^34–36^. Thus, the timing and intensity of the neuronal dynamics in each NAc cluster are likely dictated by temporally distinct glutamatergic inputs and regulated by neuromodulators and local circuits.

We identified three temporally-defined NAc neuronal clusters, among which the -5-sec and -2.5-sec clusters may correspond to key cognitive/behavioral events during the pre-press phase. One potential pre-press feature is appetitive behavior—animals are exploring and searching for reward—which relies on the proper functioning of NAc neurons^5, 37–40^. However, appetitive behaviors are typically continuous, with temporal features that do not correlate with the clear peaks of the two identified clusters. Furthermore, having been well-trained, the mice can approach rewards without major searching efforts. Rather, the -5-sec D1-neuron cluster correlated temporally with a rise in head movement without apparent locomotion initiation, suggesting changes of attention or some related cognitive events (Fig. S3K-CC). On the other hand, the -2.5-sec D2-neuron cluster correlated temporally with peak movements of the head and locomotion toward reward, suggesting the onset of approach behavior toward sucrose (Fig. S3K-CC). Together, these results suggest that the -5-sec D1-neuron cluster versus the -2.5-sec D2-neuron cluster contribute to different components of the cognitive-behavioral sequence toward lever-pressing.

During the pre-press phase, a chain of interwoven cognitive and behavioral events must occur to launch a successful operant response for sucrose. For example, the motivation for sucrose must be prioritized and engaged. In our experimental procedure, sucrose seeking and taking behaviors are self-initiated. As such, the motivation for sucrose may remain dormant (circulating in the “background”) until it is engaged. Intrinsically related to this motivation is the anticipation of reward. Occurring during the pre-press phase, anticipation of reward is correlated with increases in dopamine release in the NAc^41–44^, with such increases peaking at ∼5 sec before the operant response in rats performing self-paced sucrose seeking^45^. It is thus tempting to speculate that the -5-sec cluster preferentially contributes to the cognitive or behavioral states of motivation-engaging or reward-anticipating.

The -5-sec cluster is primarily composed of D1-neurons (Fig. 2). Extensive studies indicate a critical role for NAc D1-neurons in motivation-invigorating and reward-anticipating processes^8–11^. For example, mice establish much stronger optogenetic self-stimulation of NAc D1-neurons compared to D2-neurons, suggesting the dominant role of D1-neurons in motivation invigoration^46^. Echoing this, persistently elevating activities of NAc D1-neurons improves rewarded operant responses, while disrupting the function of NAc D1-neurons causes the opposite effects^47–50^. On the other hand, in both humans and rodents, anticipation of reward is correlated with increased NAc activity^51–53^. Extending these findings, our current study identified the temporal dynamics of D1-neurons during the pre-press phase, and demonstrated that interference with these neuronal dynamics reduced the likelihood of subsequent sucrose seeking. These results reveal that, rather than the constitutive activity of these neurons, the temporal dynamics of NAc D1-neurons are the key for engaging reward motivation to reward seeking and taking.

After motivation is engaged, mice switch from spontaneous behaviors to task performance, initiating the action sequence toward reward consumption^54^. Meanwhile, other extraneous signals must be ignored to ensure undistracted behavioral performance. These two processes are expected to occur during the late pre-press phase, for which the -2.5-sec cluster fits with respect to timeframe. While MSNs in the dorsolateral striatum have long been implicated in the initiation of rewarded action sequences^55–57^, recent results show that NAc MSNs are also essential^58, 59^. Compared to the -5-sec timepoint, we observed abrupt motion changes in mice at -2.5 sec, suggesting an initiation of new behavior (Fig. S3K-CC). The -2.5-sec cluster is mainly composed of D2-neurons, inhibition of which at -2.5 sec, but not other timepoints, decreased lever-pressing for sucrose (Fig. 4). Thus, the -2.5-sec cluster may represent another essential set of NAc dynamics that constitutes the behavioral sequence for reward seeking once the mouse is motivationally committed.

Neurons in the 0-sec cluster started increasing activities ∼1 sec before lever-pressing, which peaked upon the initiation of lever-pressing, and declined over the next 2-3 sec (Figs. 1,2). This lever-press-contingent pattern is consistent with previous findings that select populations of NAc neurons increase firing rates during the on- and post-lever-press phases^19, 60, 61^. The 0-sec cluster overlaps with at least four major motor responses, including lever-pressing, sucrose taking, sucrose consumption, and receding from sucrose magazine/sipper. In rats, Nicola et al. detect increased activities of NAc neurons during their entry, stay-in, and exit from the sucrose receptable entry^40^, suggesting motor-related functions of the 0-sec cluster. On the other hand, this cluster may also participate in the cognitive processing of unconditioned stimuli. In nonhuman primates, a population of ventral striatal neurons exhibit increased activities during rewarded operant responses, and this increase in activity is sensitive to the magnitude of the reward, but absent when only the reward-associated cues are present without reward delivery^62–65^. Furthermore, several cognitive events occur immediately after lever-pressing, such as experiencing reward and recalibrating anticipation with outcome, for which the 0-sec cluster may be involved, given the prominent role of NAc in reward learning^3, 29, 66^.

It is worth mentioning that, for most populational data, a clearcut separation of subpopulations is not possible. This is also true for our cluster analyses. For example, it appears that a small population of D1-neurons exhibited increased activities at -2.5 sec (Fig. S2KG). However, optogenetic inhibition of D1-neurons at -2.5 sec did not affect operant responding, leaving the behavioral significance of this cluster unknown. It is also worth mentioning that the three NAc clusters were identified using an operant procedure with a fixed ratio 1 schedule. Under more demanding reinforcement schedules, for which higher motivational states are required, we speculate that, if the -5-sec cluster contributes to motivation-engaging as hypothesized above, this cluster will become more prominent, e.g., exhibiting higher collective z-scores.

### D1-neurons versus D2-neurons

In D1-Cre and D2-Cre mice, Cre-dependent viral-mediated gene expression can also occur in non-MSN cell types that also express D1 or D2 receptors, although recent single cell RNA-sequencing data confirm that ∼1% of Cre+ cells are non-MSN cell types^67^. Thus, the neuronal dynamics in D1-Cre or D2-Cre mice should be interpreted with this relatively minor caveat in mind.

Increased activities of both D1- and D2-neurons are observed during reward taking, cue-conditioned reward seeking, or reward-associated learning^7, 11, 13, 68, 69^. While functionally contrasting each other on many occasions, D1- and D2-NAc neurons have also been recognized for their complementary roles in motivated behaviors^7–9, 70–72^. For example, when applied persistently in a temporally noncontingent manner, optogenetics-mediated activity downregulation of either D1- or D2-neurons decreases the likelihood of operant responding for sucrose^73^. Our results show that D1- and then D2-neurons sequentially dominate in the NAc circuit activity during the pre-press phase (Fig. 2), suggesting their distinct but temporally linked functions. Hinted by their differential coupling with motor changes (Fig. S3), we propose a heuristic model that the -5- sec D1-neuron versus -2.5-sec D2-neuron clusters preferentially contribute to distinct cognitive/behavioral events, such as motivation-engaging or sucrose-anticipating versus behavior-initiating responses. Specifically-timed inhibition of D1- or D2-neurons reduced subsequent lever-presses for sucrose (Fig. 4), indicating that both the D1-neuron and D2-neuron-dominated NAc temporal dynamics, and the cognitive/behavioral events they mediate, are essential for launching self-paced operant response for reward.

Although our current study focuses on activity increases, a portion of neurons exhibited decreased activities clustering at the three key timepoints (Figs. 1,2). These activity decreases were observed for both D1- and D2-neurons at 0 sec, such that there were indeed four clusters at 0 sec: increased D1-neurons, increased D2-neurons, decreased D1-neurons, and decreased D2-neurons (Fig. 2IJ). This observation is consistent with results from in vivo recordings that reveal distinct populations of NAc neurons exhibiting increased versus decreased activity in response to the same stimulus^19, 74, 75^. However, the decrease cluster at 0 sec is not composed of an independent neuronal population but includes a portion of the increase clusters during the pre-press phase. For example, in the two-cluster analysis of D1-neurons, most of the same neurons that exhibited increased activity at -5 sec subsequently exhibited decreased activity at ∼0 sec (Fig. 2I). Similar activity patterns were observed for D2-neurons in the -2.5-sec cluster (Fig. 2J). Thus, the decrease cluster can be regarded as the second part of a biphasic pattern of both neuronal subtypes in the -5-sec and -2.5-sec clusters. In theory, decreases and increases in neuronal activity could contribute equally to the overall circuit dynamics at a given timepoint. In this case, the increase and decrease clusters at ∼0 sec can be captured and combined by PCA, resulting in a higher separation than at ∼-5 or ∼-2.5 sec where the increase cluster alone dominated (Figs. 2IJ, 3CD). In addition, the same sets of D1- and D2-neurons participate in different functional ensembles with different temporal-dynamic features—i.e., the NAc ensemble can be organized by the timing of populational activities, not necessarily a fixed population of neurons. Furthermore, the decrease could connote inhibition, with which the corresponding cognitive or behavioral events during the pre-press phase are prevented from entering the on-press phase.

### Computational analyses

The PCA-distance results suggest that D1-versus D2-neurons enter a different activity state at ∼-5 versus ∼-2.5 sec compared to their basal activities (Figs. 3CD). In the Support Vector Machine accuracy test, the -5-sec activity state of D1-neurons and -2.5-sec activity state of D2-neurons exhibited increased correlation to subsequent lever-press actions (Fig. 3EF). Thus, both the PCA-distance and Support Vector Machine accuracy tests similarly revealed that D1- and D2-neurons exhibited temporally sequential activity states that may contribute to the pre-press cognitive/behavioral sequence.

Given the overrepresentation of pre-press neuronal activity and the ability to decode upcoming lever-presses, we also examined whether the neural trajectories through which NAc neurons reached their pre-press clustering activity states were the same, i.e., how much variability was there when NAc neurons aligned their activity to the two key pre-press timepoints? We found similar trajectories for D1- or D2-neurons (Fig. 3G-M). In fact, trajectories of D1- and D2-neurons in >75% lever-presses traveled in the same relative direction when reaching to ∼-5 or ∼-2.5 sec, respectively, suggesting that the NAc neuron-mediated processing of the pre-press cognitive-behavioral sequence is similar over trials. Extrapolated from these results are two additional conceptual considerations. First, an elevated synchronization was observed for the trajectories of D2-neurons to ∼-5 sec (Fig. S3J), the timepoint at which the activity of the -2.5-sec cluster was in the early rising phase (Figs. 1L; 2J). Thus, the trajectory synchronization may also reflect key features of NAc neurons during the ‘preparatory’ phase toward their peak activity increases. Second, at 0 sec, the activity increase of one cluster overlaps with the activity decline of another cluster (Fig. 2IJ), presumably resulting in diverse activity states and unsynchronized trajectories among neurons when clusters were sampled together. As such, although synchronized trajectories of D1 or D2-neurons were not detected at 0 sec (Fig. S3J), we speculate that such a trajectory synchronization exists in neurons from each individual cluster.

### Behavioral correlates

We speculate that the -5-sec D1-neuron cluster preferentially contributes to motivation-engaging, reward-anticipating, or other cognitive events. Our behavioral tests did not measure these cognitive events directly, but the results are in line with this speculation. For example, interfering with the -5-sec cluster activity decreased the likelihood of subsequent operant responses for sucrose (Fig. 4), an effect that may result from compromised reward motivation or anticipation. Importantly, interfering with the -5-sec cluster activity decreased the presses of both active and inactive levers, but not the ratio of active versus total (active + inactive) lever-presses (Fig. S4), suggesting that the -5-sec cluster activities are correlated with reward motivation or anticipation but not the accuracy of operant performance. In parallel, although there are basal activities of D1-neurons at -2.5 sec (Figs. 2;S2), inhibiting D1-neurons at this timepoint did not affect subsequent operant responding (Figs. 4;S4). Thus, rather than their general or constitutive activities, it is the temporally confined activities of D1-neurons that mediate key cognitive/behavioral events.

Correlated with movement changes, we speculate that the -2.5-sec cluster contributes to the initiation of the behavioral sequence toward sucrose taking. While NAc D2-neurons are often associated with behavioral inhibition, their roles in motion initiation or performance are emerging. In trial-based food taking tests, inhibition of D2-neurons immediately after predictive cues decreases the likelihood of operant responding, suggesting compromised initiation of reward seeking upon insufficient activation of D2-neurons^76^. In addition to movement initiation, NAc D2-neurons are also implicated in reward motivation. For example, mice establish intracranial self-stimulation of NAc D2-neurons, albeit with much lower magnitude compared to D1-neuron stimulation^46^. Furthermore, mice with optogenetic up-versus downregulation of NAc D2-neuron activities exhibit increased versus decreased persistence in obtaining reward, respectively^73, 77^. Thus, like D1-neurons, the behavioral function of NAc D2-neurons is also likely to be multi-dimensional, depending upon their temporal dynamics and the phases of motivational responses. Particularly for the -2.5-sec D2-neuron cluster, the sharp movement changes at -2.5 sec (Fig. S3) implicate a strong locomotor component, potentially initiating the behavioral sequence of sucrose seeking.

In summary, in reward-motivated operant responses, if the final act of lever-pressing is analogous to the takeoff of an Olympic long jump, the preluding cognitive/behavioral events would serve as the approach run. We propose that the two sets of temporal dynamics of NAc neurons identified during the pre-press phase may mediate some of these key approach-run events that launch reward-seeking behaviors. These findings thereby provide unique insight into understanding the circuit mechanisms through which motivation is engaged for behavioral output.

## Methods

### Animals, reagents, and genotyping

Wildtype C57BL/6J mice were purchased from Jackson Laboratories. Both D1R-Cre^78^ and D2R-Cre mice^78^ on a C57BL/6J background were originally purchased from Jackson Laboratories and were bred in the University of Pittsburgh animal facility. Mice were 7-weeks old at the beginning of experimentation and, after initial surgical procedures, were singly housed on a 12-hour light/dark cycle (light on/off at 7:00/19:00), at a room temperature of 22-24°C and humidity 40-60%. Mice accessed food and water ad libitum, except a few days prior to sucrose self-administration (SA) (see below). Mice were 12-14 weeks old at the initiation of behavioral experiments. Both male and female mice were used in pilot experiments, but male mice exhibited more tolerance for Miniscope wearing and stability of behavioral performance compared to female mice. Male mice were thus used for all experiments during data collection. All animal care and use were approved and performed in accordance with NIH guidelines and University of Pittsburgh’s Institutional Animal Care and Use Committee.

For optogenetic manipulations, we used Jaws-expressing adeno-associated virus (AAV) for in vivo expression, with the virus rAAV8/hsyn-Flex-Jaws-KGC-GFP-ER2 (titer ≥ 4.1×10^12^ virus molecules/ml) purchased from the University of North Carolina GTC Vector Core. For GCaMP-based in vivo Ca^2+^ imaging, we used pAAV.hsyn.GCaMP6m.WPRE.SV40 (titer ≥ 2.7 x 10^13^ GC/ml) in wildtype mice and pAAV.hsyn.Flex.GCaMP6m.WPRE.SV40 (titer ≥2.7 x 10^13^ GC/ml) in D1-Cre or D2-Cre mice, both purchased from Addgene.

Genotyping was performed using a PCR reaction containing AccuStart II GelTrack PCR SuperMix. The primers for Cre detection (5’→3’) were: forward AATGCTTCTGTCCGTTTGCC and reverse GATCCGCCGCATAACCAGT). Amplified DNA was analyzed on a 20% agarose gel with ethidium bromide staining, with the signature bands determining the genotypes^78^.

### In vivo viral injections

Mice were anesthetized with i.p. injection of ketamine (100 mg/kg)–xylazine (10 mg/kg) mixture and were held in a stereotaxic frame (Stoelting Co., Wood Dale, IL). Through a 10-μL NanoFil syringe with a 32-gauge needle controlled by UMP3 and Micro4 system (WPI, Sarasota, FL), the AAV solution was infused at a rate of 200 nL/min into the NAc (AP +1.50 mm, ML ± 0.73 mm, DV -4.25 mm). For Ca^2+^ imaging experiments, 1 µL of GCaMP6m-expressing AAV was infused unilaterally, followed by the implantation of GRIN lens (0.5 mm in diameter, 8.4 mm in length, Inscopix) directly above the viral injection site. In this process, the GRIN lens was slowly lowered into the brain tissue using a cannula holder for implantation, and a small metal bar was affixed into place via superglue and dental cement. Mice were then kept in their home cages for >4 weeks for viral expression before imaging experiments.

After >4 weeks of viral expression, the mice were anesthetized and placed in the stereotaxic frame again for installation of Miniscope baseplate. The protective cover over the GRIN lens was removed, and a Miniscope along with a relay lens (1.8 mm, 0.25 pitch, Edmund Optics) equipped with a baseplate was stereotaxically lowered above the lens. The baseplate was carefully positioned at a focal plane where fluorescent cells were visible. Once the optimal focal plane was attained, the baseplate was secured to the head with dental cement. After dental cement solidified, the Miniscope was removed, and a protective cap was secured over the GRIN lens.

### Ca^2+^ imaging in sucrose SA mice

Sucrose SA was conducted in operant chambers enclosed within sound attenuating cabinets (Med Associates). Each chamber (29.53 x 24.84 x 18.67 cm^3^) contains an active and an inactive lever, a lickometer, a sucrose sipper, a conditioned stimulus light above each lever, a house light, a speaker for audio cues, and a wide-angled infrared camera (ELP Camera USB 1080P Wide Angle Fisheye LED Infrared Webcam).

Prior to the training procedure, mice were habituated daily for >1 week (1 h/d) to head restraint, Miniscope carrying, and fiber optic cable installation. Two days prior to the overnight procedure, mice were water restricted. Following overnight training (12 h of SA session), mice underwent 11 d of self-administration training, during which pressing the active lever resulted in extraction of the sucrose sipper delivering 10% sucrose solution (0.1 mL/lick) and presentation of the compound cues (sound and light). During the 10-sec period, additional active lever-presses were recorded but not reinforced. Pressing the inactive lever did not have any consequence. The end of the 10-sec session following reward delivery was signaled by termination of the house light and retraction of the sucrose sipper. Sucrose crystalline was purchased from Fisher Chemical. Prior to each sucrose self-administration session with in vivo Ca^2+^ imaging, mice were briefly head-restrained to install the Miniscope before being placed in the operant chamber. We used a wire-free version of the UCLA Miniscope (v3)^22^. To avoid potential over-bleaching of fluorescence and best fit the capacity limit of the single cell lipo battery (Open Ephys Production), our recordings were focused on the first 20 min of each imaging session. About 40% of lever-presses occurred during the first 20 min of the 1-h SA session (raw data provided via Research Data Deposition). Thus, neuronal activities over the first 20 min may be correlated with a relatively high motivational state for sucrose.

### Ca^2+^ signal analysis

#### Preprocess by z-score standardization and down sampling

Miniscope videos of Ca^2+^ activities were recorded at a resolution of ∼320 x 320 µm and a framerate of 20 Hz. Raw video data from each imaging session were processed using an open-source package Calcium Imaging Analysis (CalmAn) in Python^79^. The CalmAn involves the motion correction, source extraction, and deconvolution steps to extract fluorescence traces of Ca^2+^ activities. Specifically, the motion artifacts of Miniscope videos were corrected by the NoRMCORRE algorithm^80^. A constrained non-negative matrix factorization (CNMF) algorithm was used to perform source extraction of fluorescence traces and eliminate overlapping spatial sources^81^. Sparse non-negative deconvolution was used to estimate the underlying neural activities^81, 82^. Following signal extraction, each trace was manually examined to exclude artifacts. To normalize signals among trials, the extracted Ca^2+^ traces were binned to 100-ms segments and z-scored with mean = 0 and standard deviation = 1. ΔF/F_0_ was used in manifold analysis, in which F_0_ was the mean z-score over 5-min sliding baseline, and ΔF = F_t_ – F_0_ at the timepoint of t.

### Heatmap and sorting

For all heatmap results, z-score data over the 20-sec time window with lever-press were extracted from individual neurons across all trials. Trial-averaged data of individual neurons were calculated by averaging the z-score data of all trials over the 20-sec time window. Data from randomly selected, trial number-matched 20-sec time windows without lever-press were used as controls. For consistent visualization, heatmaps were generated by setting the mean of trial-averaged data to 0 for individual neurons. The heatmaps were created using the *levelplot* function in the R *lattice* package (R Core Team 2022 R: A Language and Environment for Statistical Computing. R Foundation for Statistical Computing, Vienna; https://cran.r-project.org/web/packages/lattice/index.html). For sequence-based sorting heatmaps, we used two methods. The first method sorted neurons by the times at which they exhibited peak z-scores (Fig. S1CD). The second method sorted neurons by the times at which neurons exhibited initial increases in activities (Figs. 1HI). To focus on the most prominent activity changes in each neuron, we operationally grouped neurons into three sets based on their z-scores: z-scores >0.5, <0.5 but > 0.25, and <0.25, with the detection thresholds set at the z-score of 0.5, 0.25, and normalized trial-averaged z-score, respectively.

For timing-based sorting heatmaps, neurons were sorted based on normalized trial-averaged z-scores at the targeted timepoint. K-means clustering (KMC) was performed using the *kmeans* function in the R *stats* package with pre-set number of clusters to analyze the first 15 sec of the time window. For cluster presentation, we performed timing-based sorting of neurons to the cluster center, defined as the timepoints at which neurons exhibited z-scores > 95% (cutoff values) of their normalized trial-averaged z-scores during the first 15 sec of the 20-sec time window. A linear weight (= cutoff value x 4) was given to each neuron to partially filter out nonactive neurons. The cluster-wise assessment of cluster stability was performed using the *clusterboot* function in the R *fpc* package^83^.

### Collective z-scores

For collective z-score-based description of the temporal dynamics, neurons were grouped based on their KMC-allocated clusters. Activated versus nonactive neurons were defined as neurons with mean z-scores > or <0, respectively, over the ±0.5-sec range around the cluster center. Collective z-scores were calculated by summing the normalized trial-averaged z-scores of the selected neurons divided by the total number of neurons. In another grouping approach, we ranked neurons by their activity intensity around the cluster center (±0.5 sec), and included the top 50% of these neurons for subsequent plotting. The temporal changes of collective z-scores of activated neurons or top 50% of neurons ranked by activity were plotted by R package *ggplot (*https://ggplot2.tidyverse.org). Pie charts were made using R function *pie*.

### Histogram and statistics

Over the 20-sec lever-press or randomly selected trials, trial-averaged z-scores of each neuron were sliced into 0.1-sec bins. For each slice, we calculated the populational mean of z-scores by trial-averaging z-scores of all neurons. The high activity neuron was defined as the neuron with the z-score >95% of the populational mean for each bin. The counts of high activity neurons per bin were used to plot the distribution of highly active neurons over the 20-sec trials. We also examined other cutoff options (e.g., 0.3-sec bins or >90% of the populational mean) and obtained similar results.

A Chi-square test was conducted using the R function chisq.test to compare the distribution of high activity neurons between a timepoint of interest versus a randomly selected timepoint. To condense the dataset, we summed the number of high activity neurons over a 10-bin, 1-sec period across the timepoint of interest (e.g., -5.5 to -4.5 sec for the timepoint of -5 sec). We built a 2 × 2 contingency table with 2 dimensions. Dimension one was the timepoints of interest versus randomly selected timepoints with 100 repeats. Dimension two was the total counts of high activity neurons over 1 sec across the timepoints of interest versus the averaged counts of high activity neurons over each 1-sec period over the entire 20-sec trials.

#### Dimensionality Reduction

Principal component analysis (PCA) was performed to depict mathematical representation of neuronal activities. In this analysis, trial-averaged concatenated PCA was used to create a mathematical representation of neuronal activities. Activities of individual neurons over time were compared among 2 different trial types: 30-sec trials with lever-presses (15 sec before and 15 sec after the lever-press), and 30-sec random trials (baseline trials). The sample size of baseline trials was set x2 of lever-press trials to support comparison between baseline 1 and baseline 2. The mean activity of each neuron (N) over time (T) and across 2 trial types (each T x N) were concatenated into a matrix 3T x N, normalized to having mean of 0 and standard deviation of 1, and fed into the sklearn PCA model (https://scikit-learn.org/stable/about.html#citing-scikit-learn). The resulting PCs were ordered by the degrees of their explained variance. Scree plots were generated to illustrate the cumulative explained variance relative to the total variance for each PC. The first 2 PCs were selected for visualization and distance calculation (x- and y-axis in a 2-dimension Cartesian coordinate system). Thus, for a singular trial instance input T x N, we obtained the trial output instance T x 2. We also used PCA to reduce the MIND-transformed 7-dimensional manifold into 3 dimensions prior to inputting the data into the SVM. The same process of trial-averaged input to create the principal axis was used, such that we could transform a T x 7 input into a T x 3.

For PCA-related lever-press data, once we obtained their lever-press timings, we removed “overlapping” lever-presses (repeats within 10 sec from each other; e.g., for times 1, 8, 9, 15 sec we removed times 8 and 9 sec). This matched the 10-sec cooldown period in the operant procedure. We then took the filtered set of times T, and defined “lever-press” data F, so F contained all the times in the interval [t-10, t+10] for t in T. For PCA-related baseline data, we defined the baseline as all lever-press timings remaining after removing the set of lever-press intervals F. From there, we sampled |T| “lever-presses” with an available radius of 10 and defined this as our baseline sample. Note that between sampling iterations this set of “lever-presses” would change, so while it was sampled from the same original dataset of non-lever-press interval data, the data would vary slightly between runs. For MIND-related lever press data, the definition for lever-press was identical to the handling of PCA-related data. For MIND-related baseline data, we defined the baseline as time-shuffled lever presses. Thus, for an SVM testing/training on a time slice [-0.5, 0], while the lever-press data would respect its temporal relation relative to the original lever-press, baseline data could be any of the timepoints from the lever-press data F. The limitation mentioned earlier lies with how MIND generates its nonlinear transformation. Data points that lie outside of the set of lever-press data F are ill-defined (e.g., possibly null) and not suitable for a baseline definition due to the method only transforming F.

#### Centroid distance

To determine whether lever-press could be predicted by preceding neuronal activities, the spatial separation between different trial types was examined using PCA-transformed data across 20-sec time windows with or without lever-presses. The centroid of a data group was computed by calculating the mathematic mean of all data points within the group. The distance between 2 centroids [(x_1_, y_1_) and (x_2_, y_2_)] was calculated as Euclidean distance: 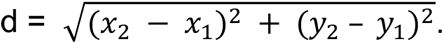 Such distances were calculated for each animal between baseline trials 1 and 2, as well as between lever-press trials and baseline trial 1, with 100 repeats (50 for pair distance) to minimize potential randomization-related biases. The group mean was computed from all animals in the group. The distance was normalized by setting the maximum distance between lever-press and baseline to “1” for each repeat.

#### Manifold inference from neural dynamics (MIND)

To generate data used for nonlinear dimensionality reduction, we smoothed ΔF/F traces from each animal with a 5-bin Gaussian filter and thresholded at 2σ, for which we measured the robust σ across the time series and individually for each neuron. We then ran the dataset through MIND^24, 84^ and embedded into 2-22 dimensions. Figure S3G shows the r^2^ values using the 7-22 dimensional embeddings of the manifold to reconstruct the raw neural activity, where r is the correlation coefficient between the raw neural activity data and the reconstructed neural activity data.

#### Support Vector Machine (SVM)

Using the MIND outputs, a series of Support Vector Machines (SVM) were built on 0.5-sec slices to determine the accuracy of neuronal activities in classifying lever-press versus baseline. An SVM was used to determine an optimal hyperplane that maximally separated different trial types. Data were split into 0.5-sec collections and then divided into a training set and a test set at the ratio of 60/40. The training set from each collection was used to train an SVM, while the test set was left out to evaluate the model’s performance. We used the sklearn LinearSVM api with the following parameters; L2 normalization penalty with squared hinge loss, regularization parameter of 0.9, no class weighting, primal optimization problem, and max iterations of 1e5. To create consistent results from randomized baseline selections, we averaged the accuracies across 100 baseline selections.

#### Bootstrapped Confidence Interval (BCI)

We bootstrapped samples and calculated the t-value of a 500-ms interval centered around the -5 and -2.5 sec to construct a 95% two-sided bootstrapped confidence interval using the scipy bootstrap and ttest_ind apis in Python. The bootstrap settings used were method=’basic’ and confidence_level=0.95. The datasets used were the rodent-type arithmetic-mean averaged set of PCA centroid distances (n = 100 samples x 5-fold points = 500 / interval) and MIND SVM prediction data (n = 100 samples / interval). The PCA centroid distance confidence interval was constructed using data centered around -5 and -2.5 (-5 ±- 0.2 sec and - 2.5 ± 0.2 sec) while the MIND data used the data from the [-5, -4.5] and [-2.5, -2.0] intervals.

### Arrow plot

Arrow plots were used to examine the directionality of the populational neural dynamics of D1- and D2-neurons in each individual mouse. Fig. 3G-L show the first 2 dimensions of manifolds embedded into 7 dimensions. Each red dot represented the time (t sec) before each lever-press, and the blue dot represented the time (t – 1.5 sec). Arrows were drawn from each blue to red dot. (t) for a timepoint of interest was determined as the time within a 1-sec range of this timepoint, at which a maximal number of arrows exhibited synchronized direction. Arrows with length <mean length were considered nonactive and excluded. In Fig. 3GIK, random 1.5-sec intervals with the number matching lever-press trials were similarly selected, aligned, and plotted. Arrows within 150° (±75°) of the mean direction (averaged from all arrows) were regarded as having synchronized direction; they were colored in blue and represent “synced angles”. All other arrows were colored in red. The proportion of arrows classified as “synced angles” is shown in Figure 3M and Figure S3J.

### Slice electrophysiology

To prepare acute brain slices^85^, mice were decapitated under isoflurane anesthesia. Coronal slices (250-μm thick) containing the NAc were prepared on a VT1200S vibratome (Leica) in a 4°C cutting solution containing (in mM): 135 N-methyl-d-glucamine, 1 KCl, 1.2 KH_2_PO_4_, 0.5 CaCl_2_, 1.5 MgCl_2_, 20 choline-HCO_3_, and 11 glucose, saturated with 95% O_2_ / 5% CO_2_, and pH adjusted to 7.4 with HCl. Osmolality of cutting solution was adjusted to 305-309 mOsm. Slices were incubated in the artificial cerebrospinal fluid (aCSF) containing (in mM): 119 NaCl, 2.5 KCl, 2.5 CaCl_2_, 1.3 MgCl_2_, 1 NaH_2_PO_4_, 26.2 NaHCO_3_, and 11 glucose, with the osmolality adjusted to 290-295 mOsm and saturated with 95% O_2_ / 5% CO_2_. The brain slices were incubated at 34°C for 30 min and then allowed to recover for >30 min at 20-22°C before electrophysiological recordings.

### Optogenetic electrophysiology

As detailed in previous studies^86–89^, current-clamp whole-cell recordings on NAc neurons were performed using an Axon MultiClamp 700B amplifier (Molecular Devices, San Jose, CA). Borosilicate glass pipettes pulled on a P-97 Puller (Sutter Instruments, Novato, CA), with the resistance of 2-5 MΩ when filled with a potassium-based internal solution containing (in mM): 130 K-methanesulfate, 10 KCl, 10 Hepes, 0.4 EGTA, 2 MgCl_2_, 3 Mg-ATP, 0.25 Na-GTP; pH 7.3 and osmolarity adjusted to 285-290 mOsm. Jaws-expressing versus noninfected control neurons were differentiated by the presence versus absence of GFP fluorescence. Action potentials were evoked by current steps multiple times with or without optogenetic stimulation (250 ms, 635-nm laser, 1-2 mW through fluorescent port of the microscope; Sloc Lasers). The timing and duration of laser stimulation as well as the recording were controlled by preprogrammed Clampex 9.2 (Molecular Devices). Data were on-line filtered at 2.6-3 kHz, amplified 5 times, and digitized at 20 kHz.

### Time-specific interference of neuronal clusters

#### In vivo virus injection and implantation of optical fiber

To perform in vivo laser stimulation of Jaws-expressing D1- or D2-neurons, D1- or D2-Cre mice were injected bilaterally with rAAV8/hsyn-Flex-Jaws-KGC-GFP-ER2 (UNC GTC Vector Core) into the NAc (in mm: AP +1.50; ML ± 0.73; DV -4.25). After stereotaxic insertion of injection needles, infusion of virus was started at a rate of 200 nL/min. After infusion, the needle was held in place for 5 min before removal to minimize the backflow of viral solution. After viral injection, the dual cannulae (0.37 NA, 1.5 mm Pitch, Ø200 µm, Doric Lenses Inc. Quebec, QC, Canada) within a guiding socket receptacle were bilaterally inserted into the NAc (in mm: AP 1.50; ML ±1.8; DV −4.2). Dental cement and screws were used to secure the cannulae on the skull.

#### In vivo optogenetics

Laser pulses were applied bilaterally to the NAc in freely moving mice (8 D1-Cre and 7 D2-Cre mice) via Ø200-µm optical fibers connected to a Splitter Branching Patchcord (Core: 200 µm, NA: 0.37, Jacket: 900 µm; Doric Lenses). Prior to each sucrose SA session with in vivo optogenetics, mice were head-restrained and the patchcord was connected to optical fibers. The patchcord was connected to a 635-nm laser diode controller (Sloc Lasers), and laser pulses (pulse duration, 250 ms) were generated through a waveform generator (Master 8). The light intensity through the patchcord was measured by a light sensor (S130A, Thor Labs) and adjusted to 5-10 mW prior to behavioral sessions. We trained the DLC algorithms for each mouse during the “off” session and then perform the test during an “on” session on the next day for a timepoint. We then trained the mice for two “off” sessions over another two days, to monitor behavioral stability and retrain the DLC. We then started another “on” session on the following day for another timepoint. Over this procedure, the mean performance of mice was similar over the “off” sessions, suggesting stable behavioral output under the control condition. The behavioral procedure for these optogenetics experiments was identical to the experiments for vivo imaging, except laser stimulation.

### DeepLabCut (DLC)

We used DLC to label individual body parts, which were then converted to digital coordinates. The DLC algorithm used the sequences of movements in a long short-term memory (LSTM) model to predict lever-press in real time with low-latency^27^. Once a lever-press was predicted, the model sent an output with a custom code in Arduino (Uno R3) to the laser generator, which delivered a 250-ms laser stimulation via the patchcord at specified timepoints.

#### DLC-based Prediction Model

DLC model^90, 91^ was trained using a complied 10-h sucrose SA video, in which we manually labeled 10 body and equipment parts. Key parameters for training were configured as: augmenter_type was set to ‘imgaug,’ and the ImageNet pre-trained network used was ResNet-152. Other parameters were maintained at the DLC default settings. During training with optogenetic manipulations, the coordinates of 10 moving points were generated using DLC-live through the chamber camera. The DLC-live recorded videos had a frame rate ranging from 25 to 33 frames/sec. Lever-presses during the timeout period and those that were partially overlapped were filtered out. The coordinates were grouped by 31 frames (∼1 sec) / segment. Coordinate segments before lever-presses were used as positive samples. Coordinate segments of randomly selected timeframes excluding positive samples were used as negative samples. For the positive samples in the -5-, -2.5-, and -0.5-sec models, we used the data from -6 to -5 sec, -4 to -3 sec, and -2.5 to -1.5 sec, respectively. Data augmentation was implemented by shifting ∼1 sec of frames backward or forward, resulting in a x29-fold augmentation of the size of positive samples. The size of negative samples was determined in two ways: 1) for mice with the number of verified active lever-presses >150 (∼averaged number of lever-presses), the size of negative samples was adjusted to match the size of positive samples; 2) for mice with the number of active-presses <150 active lever-presses, the size of negative samples was set to match 150 lever-presses. These samples were then fitted to a PyTorch-constructed LSTM model (arXiv:1912.01703) with hidden dimensions set to 256, layer dimensions to 8, and a learning rate of 0.0001 using the torch.nn.LSTM function. Randomized predictions were made by a random model, which generated random positive versus negative outputs using the positive-negative frequencies determined by the input samples of each mouse. A filter was applied to the LSTM such that the positive outputs must repeat >1 over 1 sec to generate a positive prediction.

The balanced accuracy of DLC prediction was calculated using the formula: (true positives / positive samples + true negatives / negative samples) / 2. The Positive Predictive Value (PPV) was calculated as: true positives / predicted positives. Relative PPV in each mouse was calculated as: PPV (on) / PPV (off). All lever-presses were treated as positive samples. Correction predictions of lever-presses were considered a true positive. Negative samples in the balanced accuracy and PPV calculation were collected at a rate of one sample / sec.

#### DLC-based calculation of movement

The velocity of a mouse’s movement was calculated based on 10 DLC labels (‘nose’, ‘objectA’, ‘left ear’, ‘right ear’, ‘neck’, ‘middle back’, ‘middle left’, ‘middle right’, ‘tail bottom’, ‘tail mid’) that traced the coordinates of different body parts. We categorized the labels into three parts: head (‘nose’, ‘objectA’, ‘left ear’, ‘right ear’), body (‘neck’, ‘middle back’, ‘middle left’, ‘middle right’), and tail (‘tail bottom’, ‘tail mid’). For both D1 and D2, we performed the calculation on 7 animals, using the day 9 and day 11 data, across a total of 14 experiments. We determined the trial-averaged speed for each body part in two types of trials. One type involved 20-sec trials with lever-presses (from 10 sec before to 10 sec after lever-pressing), and the other consisted of 20-sec random trials (baseline trials). To compute the speed at each timepoint, we filtered out coordinates with low confidence values (<0.95 for head and body, <0.5 for tail). We then measured the Euclidean distance between the coordinates of the current timepoint (x1, y1) and the next timepoint (x2, y2), which was 0.1 sec after the current timepoint. After that we divided the distance by the time elapsed between them (0.1s) to calculate the averaged movement velocity of each body part. 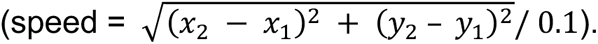 The trial-averaged velocity was obtained from individual animals for statistical summarization, including the mean and the standard error. The plots were generated by the Python package Seaborn.

#### Regression

To explore the correlation between the movement and neuronal activity, we selected 20-sec trials with lever-presses (from 10 sec before to 10 sec after lever-pressing) and calculated the trial-averaged velocity as well as trial-averaged neuronal activity. Then we put velocity as input and neuronal activity as output and used function stats.linregress in Python package scipy to fit a linear regression. We used the Pearson’s correlation coefficient to assess the relationship between velocity and neuronal activity. To analyze such correlations at -5, -2.5 and -0.5 sec, we chose the data over the intervals of -5.5 to -4.5 sec, -3 to -2 sec and -0.5 to 0.5 sec, and calculated the orthogonal distance between the datapoints in each time interval and the regression line 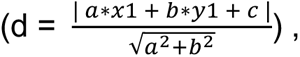 where the point is (x1, y1) and the line is *a* ∗ *x* + *b* ∗ *y* + *c* = 0. The regression plots were generated using the Python package Matplotlib. The statistical figures were generated using GraphPad Prism.

#### Location

The location of a mouse at a given timepoint was calculated using the mean coordinates of the head part of the labels (‘nose’, ‘objectA’, ‘left ear’, ‘right ear’). We also calculated the mean and the standard deviation of all individual locations. Both individual location plots and mean location plots were generated using the Python package Matplotlib. The distance between the location of each animal at each timepoint (x1, y1) and the mean location at -0.5s (x2, y2) was calculated using Euclidean distance 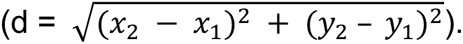 The statistical figures were generated using GraphPad Prism.

#### Direction

The direction of a mouse from one timepoint to another was calculated using atan2 function and degrees function in Python package Math. The arrow plots were generated using the Python package Matplotlib. For the statistical figures, we first calculated the angle from the starting timepoint location towards the active lever, as well as the angle from the starting timepoint location towards the sipper. Then we subtracted the two values by the angle from the starting timepoint location towards next timepoint location, and selected the minimum of the absolute values of these two as the angle difference. We performed these calculations for all pairs of locations. The statistical figures were generated using GraphPad Prism.

### Staining and confocal imaging

After deep anesthesia with isoflurane, mice were perfused transcardially first with 20 mL of 0.01 M phosphate-buffered saline (PBS, pH 7.4) and then with 20 mL of 4% paraformaldehyde in 0.1 M phosphate buffer. The brains were removed and postfixed in 4% paraformaldehyde at 4°C for 6-8 h, followed by incubation with sucrose solutions of graded concentrations (20% then 30%) in 0.01 M PBS at 4°C for 24 h. The brain was sectioned into 35 µm-thick slices at -20°C using a cryostat microtome (CryoStar NX50, ThermoFisher). The brain slices containing the NAc (+1.8 to 0.8 mm from bregma) were washed by 0.01 M PBS, then blocked for 1 h in 4% normal goat serum and 0.4% Triton X-100 (TBS) in 0.01M PBS. The sections were then incubated with the primary antibody (rabbit anti-Cre, 1:300 diluted in TBS, Cell Signaling Technology, #15036) for 48 h at 4°C with gentle shaking. Next, the sections were rinsed in 0.01 M PBS for 3 times (5 min each time) and then incubated with the secondary antibody (goat anti-rabbit Alexa Fluor 568, 1:150 diluted in TBS ThermoFisher, #A11011) for 2 h. The sections were then rinsed in 0.01 M PBS for 3 times (5 min each time). After rinsing, the sections were mounted on glass slides with Mounting Medium with DAPI (Abcam, #AB104139). Fluorescence images were captured with a Leica TCS SP8 confocal laser-scanning microscope, using a 10X lens for low magnification and a 63X oil immersion objective for high magnification.

### Data acquisition and statistics

For in vivo Ca^2+^ imaging experiments, data were collected from 7 wildtype, 5 D1-Cre, and 6 D2-cre mice. For optogenetic electrophysiology, data were collected from 5-6 brain slices from 3 mice. For in vivo optogenetics experiments, data were collected from 8 D1-Cre and 7 D2-Cre mice. Data of the optogenetic inhibition at -0.5 sec were unable to collected from one D1-Cre mouse, due to unexpected incident (the mouse jumped out of the test chamber, resulting in termination of the experiment). Unaffected portions of data from this mouse were still used for DeepLabCut training. No statistical methods were used to predetermine sample sizes, but our sample sizes were consistent with those reported in previous publications with similar experimental designs^73, 92–94^. All data collection was randomized. All data were analyzed offline, and investigators were not blinded to experimental conditions during the analyses. Statistical analyses were performed in GraphPad Prism (v10) or R (v4). All data collected from verified procedures were included in the final statistical analysis. Statistical significance was assessed using paired t-test or two-tailed one-way ANOVA repeated measures, followed by Bonferroni’s posttests. Differences were considered significant when the p value <0.05. Statistical results were expressed as mean ± s.e.m. Standard deviations were used for some figure presentations.

## Data Availability

Source data are reported in Source Data sheet published together with this manuscript.

## Code Availability

All custom codes can be found at Code Ocean through the link: https://codeocean.com/capsule/0290769/tree/v1.

## Acknowledgements

We thank Min Li for excellent technical support, and Drs. Yavin Shaham and Terry Robinson for suggestions on behavioral models. The authors’ work was partially supported by NIH grants DA053388 (EHN), DA023206 (YD), DA060868 (YD), and DA040620 (EJN,YD).

## Author Contributions

Conceptualization: TAS, K-LL, XQ, WJW, YHH, OMS, EJN, EJN, YD; Experimentation: TAS, K-LL, WJW, EEA, RJZ, QL; Data Analysis: TAS, K-LL, XQ, BL, JL, QL, BZ, LW, EHN, YD; Data Interpretation: all authors; Manuscript writing: TAS, K-LL, XQ, BL, WJW, YHH, EJN, EHN, YD.

---

**Figure S1.**
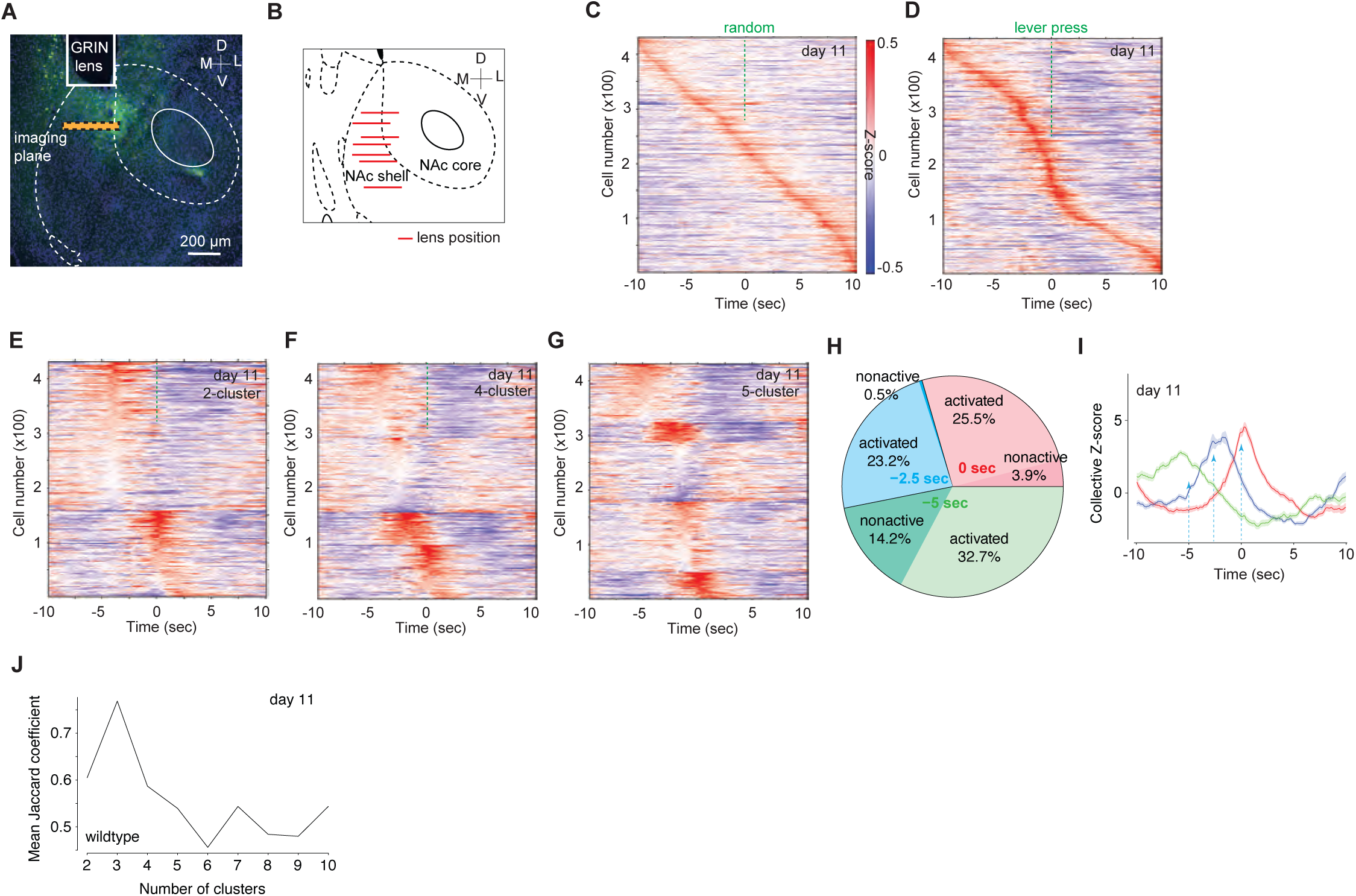

**Figure S2.**
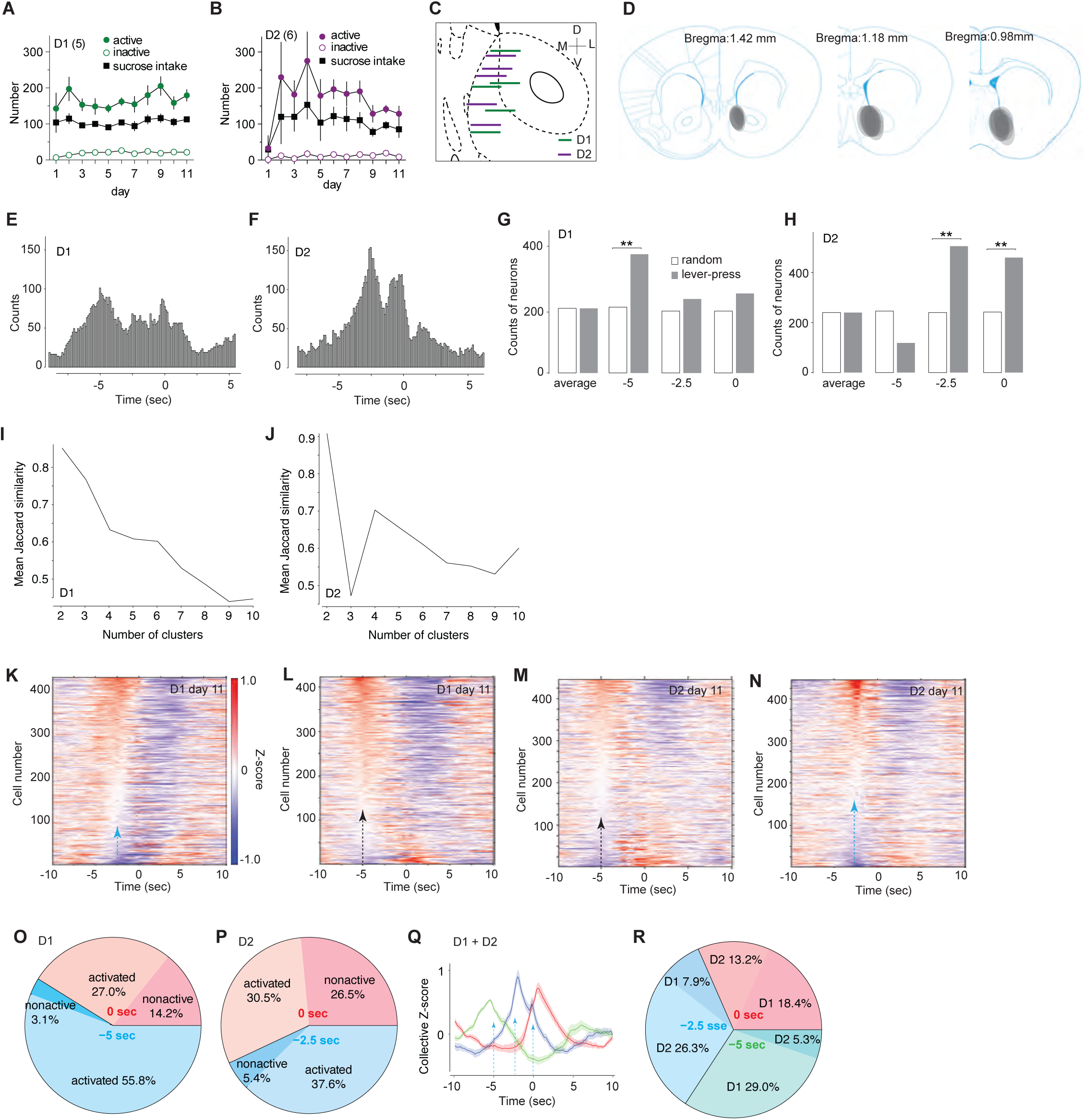

**Figure S3.**
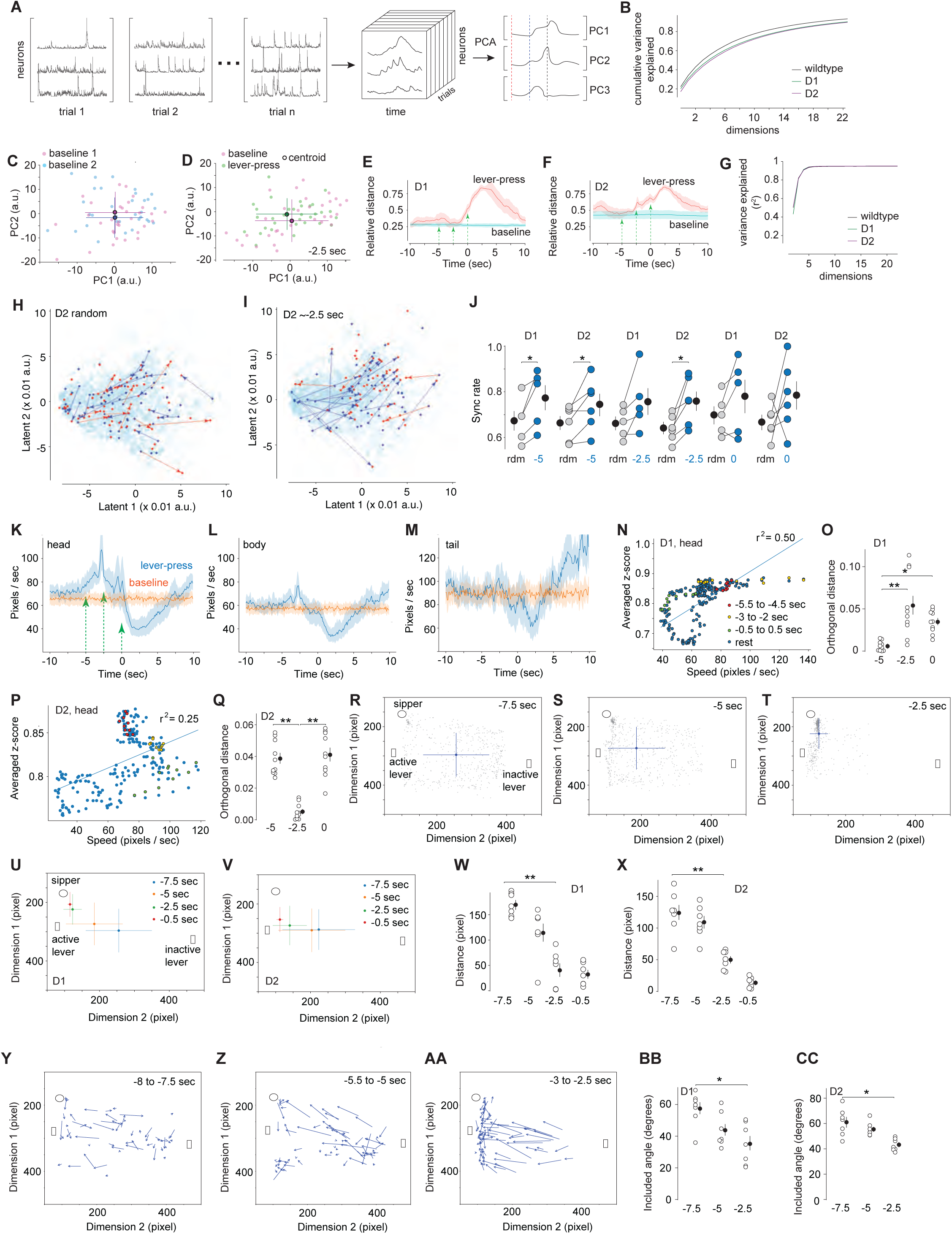

**Figure S4.**
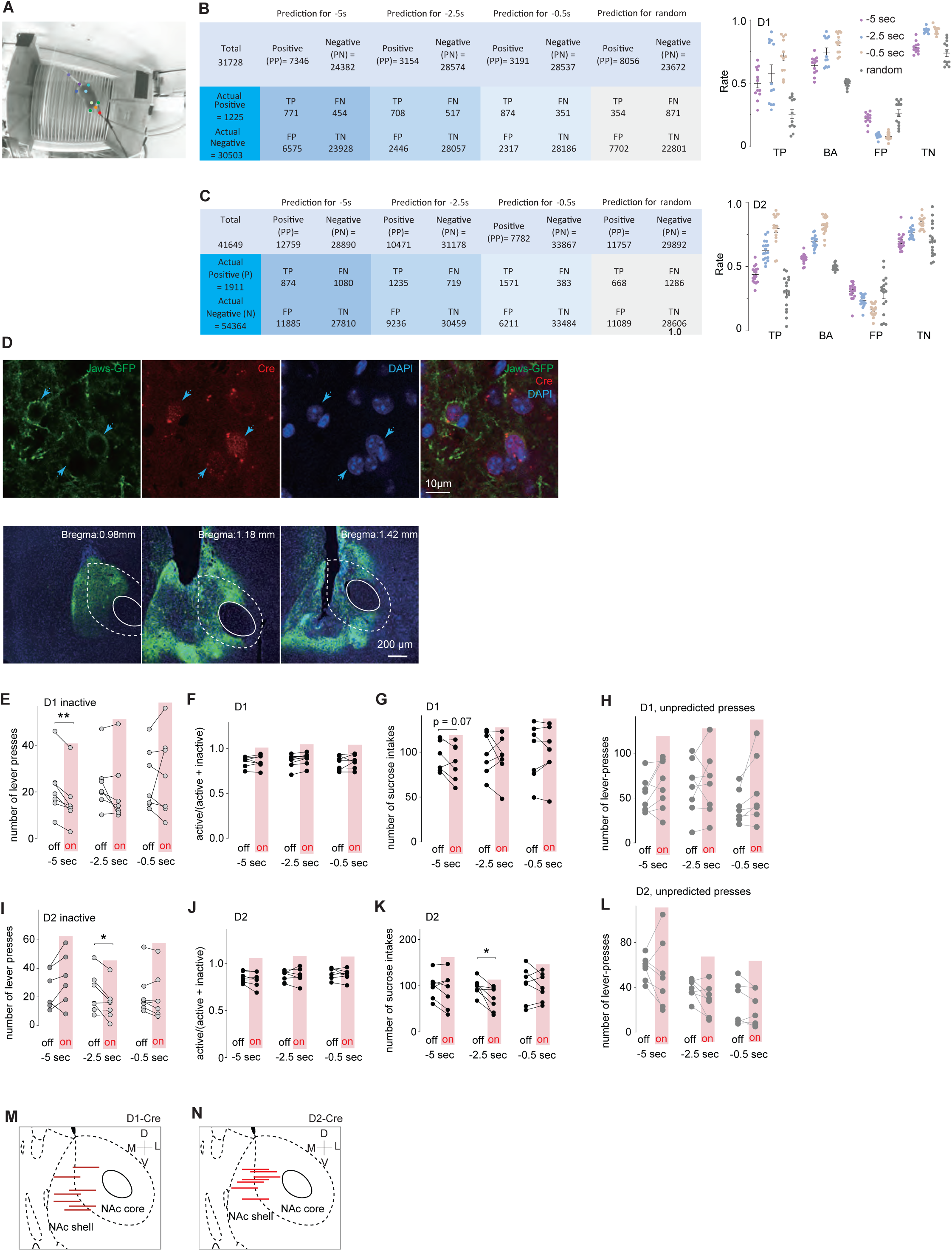

## References

1. Swanson, L.W. The neural basis of motivated behavior. Acta Morphol Neerl Scand 26, 165–176 (1988).

2. Puglisi-Allegra, S. & Ventura, R. Prefrontal/accumbal catecholamine system processes emotionally driven attribution of motivational salience. Rev Neurosci 23, 509–526 (2012).

3. Yin, H.H., Ostlund, S.B. & Balleine, B.W. Reward-guided learning beyond dopamine in the nucleus accumbens: the integrative functions of cortico-basal ganglia networks. The European journal of neuroscience 28, 1437–1448 (2008).

4. Cardinal, R.N., Parkinson, J.A., Hall, J. & Everitt, B.J. Emotion and motivation: the role of the amygdala, ventral striatum, and prefrontal cortex. Neurosci Biobehav Rev 26, 321–352 (2002).

5. Berridge, K.C. & Kringelbach, M.L. Pleasure systems in the brain. Neuron 86, 646–664 (2015).

6. Gong, S., et al. A gene expression atlas of the central nervous system based on bacterial artificial chromosomes. Nature 425, 917–925 (2003).

7. Allichon, M.C., et al. Cell-Type-Specific Adaptions in Striatal Medium-Sized Spiny Neurons and Their Roles in Behavioral Responses to Drugs of Abuse. Front Synaptic Neurosci 13, 799274 (2021).

8. Zinsmaier, A.K., Dong, Y. & Huang, Y.H. Cocaine-induced projection-specific and cell type-specific adaptations in the nucleus accumbens. Mol Psychiatry 27, 669–686 (2022).

9. Lobo, M.K. & Nestler, E.J. The striatal balancing act in drug addiction: distinct roles of direct and indirect pathway medium spiny neurons. Front Neuroanat 5, 41 (2011).

10. Smith, R.J., Lobo, M.K., Spencer, S. & Kalivas, P.W. Cocaine-induced adaptations in D1 and D2 accumbens projection neurons (a dichotomy not necessarily synonymous with direct and indirect pathways). Curr Opin Neurobiol 23, 546–552 (2013).

11. Calipari, E.S., et al. In vivo imaging identifies temporal signature of D1 and D2 medium spiny neurons in cocaine reward. Proc Natl Acad Sci U S A 113, 2726–2731 (2016).

12. Tan, B., et al. Drugs of abuse hijack a mesolimbic pathway that processes homeostatic need. Science 384, eadk6742 (2024).

13. Tan, B., et al. Dynamic processing of hunger and thirst by common mesolimbic neural ensembles. Proc Natl Acad Sci U S A 119, e2211688119 (2022).

14. Pennartz, C.M., Groenewegen, H.J. & Lopes da Silva, F.H. The nucleus accumbens as a complex of functionally distinct neuronal ensembles: an integration of behavioural, electrophysiological and anatomical data. Prog Neurobiol 42, 719–761 (1994).

15. Deadwyler, S.A., Hayashizaki, S., Cheer, J. & Hampson, R.E. Reward, memory and substance abuse: functional neuronal circuits in the nucleus accumbens. Neurosci Biobehav Rev 27, 703–711 (2004).

16. Wright, W.J. & Dong, Y. Silent Synapses in Cocaine-Associated Memory and Beyond. J Neurosci 41, 9275–9285 (2021).

17. Salery, M., et al. Transcriptional correlates of cocaine-associated learning in striatal ARC ensembles. bioRxiv (2023).

18. Day, J.J., Jones, J.L. & Carelli, R.M. Nucleus accumbens neurons encode predicted and ongoing reward costs in rats. The European journal of neuroscience 33, 308–321 (2011).

19. Roitman, M.F., Wheeler, R.A. & Carelli, R.M. Nucleus accumbens neurons are innately tuned for rewarding and aversive taste stimuli, encode their predictors, and are linked to motor output. Neuron 45, 587–597 (2005).

20. Atallah, H.E., McCool, A.D., Howe, M.W. & Graybiel, A.M. Neurons in the ventral striatum exhibit cell-type-specific representations of outcome during learning. Neuron 82, 1145–1156 (2014).

21. Gill, T.M., Castaneda, P.J. & Janak, P.H. Dissociable roles of the medial prefrontal cortex and nucleus accumbens core in goal-directed actions for differential reward magnitude. Cereb Cortex 20, 2884–2899 (2010).

22. Cai, D.J., et al. A shared neural ensemble links distinct contextual memories encoded close in time. Nature 534, 115–118 (2016).

23. Li, Y., et al. Neuronal Representation of Social Information in the Medial Amygdala of Awake Behaving Mice. Cell 171, 1176–1190 e1117 (2017).

24. Nieh, E.H., et al. Geometry of abstract learned knowledge in the hippocampus. Nature 595, 80–84 (2021).

25. Cortes, C. & Vapnik, V. Support-vector networks. Mach Learn 20, 273–297 (1995).

26. Baker, M., et al. External globus pallidus input to the dorsal striatum regulates habitual seeking behavior in male mice. Nat Commun 14, 4085 (2023).

27. Kane, G.A., Lopes, G., Saunders, J.L., Mathis, A. & Mathis, M.W. Real-time, low-latency closed-loop feedback using markerless posture tracking. Elife 9 (2020).

28. Chuong, A.S., et al. Noninvasive optical inhibition with a red-shifted microbial rhodopsin. Nat Neurosci 17, 1123–1129 (2014).

29. Wright, W.J. & Dong, Y. Psychostimulant-Induced Adaptations in Nucleus Accumbens Glutamatergic Transmission. Cold Spring Harb Perspect Med 10 (2020).

30. Yu, J., et al. Contingent Amygdala Inputs Trigger Heterosynaptic LTP at Hippocampus-to-Accumbens Synapses. J Neurosci (2022).

31. Xia, S.H., et al. Cortical and Thalamic Interaction with Amygdala-to-Accumbens Synapses. J Neurosci 40, 7119–7132 (2020).

32. Ishikawa, M., et al. Dopamine Triggers Heterosynaptic Plasticity. Journal of Neuroscience 33, 6759–6765 (2013).

33. Yu, J., et al. Ventral Tegmental Area Projection Regulates Glutamatergic Transmission in Nucleus Accumbens. Sci Rep 9, 18451 (2019).

34. Schall, T.A., Wright, W.J. & Dong, Y. Nucleus accumbens fast-spiking interneurons in motivational and addictive behaviors. Mol Psychiatry 26, 234–246 (2021).

35. Yu, J., et al. Nucleus accumbens feedforward inhibition circuit promotes cocaine self-administration. Proc Natl Acad Sci U S A 114, E8750–E8759 (2017).

36. Wright, W.J., Schluter, O.M. & Dong, Y. A Feedforward Inhibitory Circuit Mediated by CB1-Expressing Fast-Spiking Interneurons in the Nucleus Accumbens. Neuropsychopharmacology 42, 1146–1156 (2017).

37. Berridge, K.C. Food reward: brain substrates of wanting and liking. Neurosci Biobehav Rev 20, 1–25 (1996).

38. Nicola, S.M. Reassessing wanting and liking in the study of mesolimbic influence on food intake. Am J Physiol Regul Integr Comp Physiol 311, R811–R840 (2016).

39. Taha, S.A. & Fields, H.L. Encoding of palatability and appetitive behaviors by distinct neuronal populations in the nucleus accumbens. J Neurosci 25, 1193–1202 (2005).

40. Nicola, S.M., Yun, I.A., Wakabayashi, K.T. & Fields, H.L. Firing of nucleus accumbens neurons during the consummatory phase of a discriminative stimulus task depends on previous reward predictive cues. J Neurophysiol 91, 1866–1882 (2004).

41. Goedhoop, J., Arbab, T. & Willuhn, I. Anticipation of Appetitive Operant Action Induces Sustained Dopamine Release in the Nucleus Accumbens. J Neurosci 43, 3922–3932 (2023).

42. Wakabayashi, K.T., Fields, H.L. & Nicola, S.M. Dissociation of the role of nucleus accumbens dopamine in responding to reward-predictive cues and waiting for reward. Behav Brain Res 154, 19–30 (2004).

43. Saddoris, M.P., Cacciapaglia, F., Wightman, R.M. & Carelli, R.M. Differential Dopamine Release Dynamics in the Nucleus Accumbens Core and Shell Reveal Complementary Signals for Error Prediction and Incentive Motivation. J Neurosci 35, 11572–11582 (2015).

44. Saddoris, M.P., Sugam, J.A., Cacciapaglia, F. & Carelli, R.M. Rapid dopamine dynamics in the accumbens core and shell: learning and action. Front Biosci (Elite Ed) 5, 273–288 (2013).

45. Wassum, K.M., Ostlund, S.B. & Maidment, N.T. Phasic mesolimbic dopamine signaling precedes and predicts performance of a self-initiated action sequence task. Biol Psychiatry 71, 846–854 (2012).

46. Cole, S.L., Robinson, M.J.F. & Berridge, K.C. Optogenetic self-stimulation in the nucleus accumbens: D1 reward versus D2 ambivalence. PLoS One 13, e0207694 (2018).

47. Bin Saifullah, M.A., et al. Cell type-specific activation of mitogen-activated protein kinase in D1 receptor-expressing neurons of the nucleus accumbens potentiates stimulus-reward learning in mice. Sci Rep 8, 14413 (2018).

48. Macpherson, T. & Hikida, T. Nucleus Accumbens Dopamine D1-Receptor-Expressing Neurons Control the Acquisition of Sign-Tracking to Conditioned Cues in Mice. Front Neurosci 12, 418 (2018).

49. Matikainen-Ankney, B.A., et al. Nucleus Accumbens D(1) Receptor-Expressing Spiny Projection Neurons Control Food Motivation and Obesity. Biol Psychiatry 93, 512–523 (2023).

50. Derman, R.C., Bryda, E.C. & Ferrario, C.R. Role of nucleus accumbens D1-type medium spiny neurons in the expression and extinction of sign-tracking. Behav Brain Res 459, 114768 (2024).

51. Cohen, M.X., et al. Top-down-directed synchrony from medial frontal cortex to nucleus accumbens during reward anticipation. Hum Brain Mapp 33, 246–252 (2012).

52. Miyazaki, K., Mogi, E., Araki, N. & Matsumoto, G. Reward-quality dependent anticipation in rat nucleus accumbens. Neuroreport 9, 3943–3948 (1998).

53. Martin, P.D. & Ono, T. Effects of reward anticipation, reward presentation, and spatial parameters on the firing of single neurons recorded in the subiculum and nucleus accumbens of freely moving rats. Behav Brain Res 116, 23–38 (2000).

54. Nicola, S.M. The flexible approach hypothesis: unification of effort and cue-responding hypotheses for the role of nucleus accumbens dopamine in the activation of reward-seeking behavior. J Neurosci 30, 16585–16600 (2010).

55. Masimore, B., Schmitzer-Torbert, N.C., Kakalios, J. & Redish, A.D. Transient striatal gamma local field potentials signal movement initiation in rats. Neuroreport 16, 2021–2024 (2005).

56. Jin, X. & Costa, R.M. Start/stop signals emerge in nigrostriatal circuits during sequence learning. Nature 466, 457–462 (2010).

57. Jog, M.S., Kubota, Y., Connolly, C.I., Hillegaart, V. & Graybiel, A.M. Building neural representations of habits. Science 286, 1745–1749 (1999).

58. Fraser, K.M., Chen, B.J. & Janak, P.H. Nucleus accumbens and dorsal medial striatal dopamine and neural activity are essential for action sequence performance. The European journal of neuroscience (2023).

59. Collins, A.L., et al. Dynamic mesolimbic dopamine signaling during action sequence learning and expectation violation. Sci Rep 6, 20231 (2016).

60. Gutman, A.L. & Taha, S.A. Acute ethanol effects on neural encoding of reward size and delay in the nucleus accumbens. J Neurophysiol 116, 1175–1188 (2016).

61. Shidara, M. & Richmond, B.J. Differential encoding of information about progress through multi-trial reward schedules by three groups of ventral striatal neurons. Neurosci Res 49, 307–314 (2004).

62. Cromwell, H.C. & Schultz, W. Effects of expectations for different reward magnitudes on neuronal activity in primate striatum. J Neurophysiol 89, 2823–2838 (2003).

63. Hassani, O.K., Cromwell, H.C. & Schultz, W. Influence of expectation of different rewards on behavior-related neuronal activity in the striatum. J Neurophysiol 85, 2477–2489 (2001).

64. Apicella, P., Legallet, E. & Trouche, E. Responses of tonically discharging neurons in the monkey striatum to primary rewards delivered during different behavioral states. Exp Brain Res 116, 456–466 (1997).

65. Hollerman, J.R., Tremblay, L. & Schultz, W. Influence of reward expectation on behavior-related neuronal activity in primate striatum. J Neurophysiol 80, 947–963 (1998).

66. Corbit, L.H. & Balleine, B.W. Learning and Motivational Processes Contributing to Pavlovian-Instrumental Transfer and Their Neural Bases: Dopamine and Beyond. Curr Top Behav Neurosci 27, 259–289 (2016).

67. Mews, P., Mason, A.V., Kirchner, E.G., Estill, M. & Nestler, E.J. Cocaine-Induced Gene Regulation in D1 and D2 Neuronal Ensembles of the Nucleus Accumbens Revealed by Single-Cell RNA Sequencing. bioRxiv, 2024.2006.2019.599762 (2024).

68. Zachry, J.E., et al. D1 and D2 medium spiny neurons in the nucleus accumbens core have distinct and valence-independent roles in learning. Neuron (2023).

69. Tan, B., et al. Drugs of abuse hijack a mesolimbic pathway that processes homeostatic need. bioRxiv, 2023.2009.2003.556059 (2023).

70. Domingues, A.V., Rodrigues, A.J. & Soares-Cunha, C. A novel perspective on the role of nucleus accumbens neurons in encoding associative learning. FEBS Lett 597, 2601–2610 (2023).

71. Varin, C., Cornil, A., Houtteman, D., Bonnavion, P. & de Kerchove d’Exaerde, A. The respective activation and silencing of striatal direct and indirect pathway neurons support behavior encoding. Nat Commun 14, 4982 (2023).

72. Walle, R., et al. Nucleus accumbens D1- and D2-expressing neurons control the balance between feeding and activity-mediated energy expenditure. Nat Commun 15, 2543 (2024).

73. Soares-Cunha, C., et al. Activation of D2 dopamine receptor-expressing neurons in the nucleus accumbens increases motivation. Nat Commun 7, 11829 (2016).

74. Roop, R.G., Hollander, J.A. & Carelli, R.M. Accumbens activity during a multiple schedule for water and sucrose reinforcement in rats. Synapse 43, 223–226 (2002).

75. Roitman, M.F., Wheeler, R.A., Wightman, R.M. & Carelli, R.M. Real-time chemical responses in the nucleus accumbens differentiate rewarding and aversive stimuli. Nat Neurosci 11, 1376–1377 (2008).

76. Natsubori, A., et al. Ventrolateral Striatal Medium Spiny Neurons Positively Regulate Food-Incentive, Goal-Directed Behavior Independently of D1 and D2 Selectivity. J Neurosci 37, 2723–2733 (2017).

77. Soares-Cunha, C., et al. Distinct role of nucleus accumbens D2-MSN projections to ventral pallidum in different phases of motivated behavior. Cell Rep 38, 110380 (2022).

78. Gong, S., et al. Targeting Cre recombinase to specific neuron populations with bacterial artificial chromosome constructs. J Neurosci 27, 9817–9823 (2007).

79. Giovannucci, A., et al. CaImAn an open source tool for scalable calcium imaging data analysis. Elife 8 (2019).

80. Pnevmatikakis, E.A. & Giovannucci, A. NoRMCorre: An online algorithm for piecewise rigid motion correction of calcium imaging data. J Neurosci Methods 291, 83–94 (2017).

81. Pnevmatikakis, E.A., et al. Simultaneous Denoising, Deconvolution, and Demixing of Calcium Imaging Data. Neuron 89, 285–299 (2016).

82. Vogelstein, J.T., et al. Fast nonnegative deconvolution for spike train inference from population calcium imaging. J Neurophysiol 104, 3691–3704 (2010).

83. Hennig, C. Cluster-wise assessment of cluster stability. Comput Stat Data An 52, 258–271 (2007).

84. Low, R.J., Lewallen, S., Aronov, D., Nevers, R. & Tank, D.W. Probing variability in a cognitive map using manifold inference from neural dynamics. bioRxiv, 418939 (2018).

85. Neumann, P.A., et al. Increased excitability of lateral habenula neurons in adolescent rats following cocaine self-administration. Int J Neuropsychopharmacol 18 (2014).

86. Hu, X.T., Dong, Y., Zhang, X.F. & White, F.J. Dopamine D2 receptor-activated Ca2+ signaling modulates voltage-sensitive sodium currents in rat nucleus accumbens neurons. J Neurophysiol 93, 1406–1417 (2005).

87. He, Y., et al. Membrane excitability of nucleus accumbens neurons gates the incubation of cocaine craving. Neuropsychopharmacology 48, 1318–1327 (2023).

88. Wang, J., et al. Cascades of Homeostatic Dysregulation Promote Incubation of Cocaine Craving. J Neurosci 38, 4316–4328 (2018).

89. Shukla, A., et al. Calcium-permeable AMPA receptors and silent synapses in cocaine-conditioned place preference. EMBO J 36, 458–474 (2017).

90. Mathis, A., et al. DeepLabCut: markerless pose estimation of user-defined body parts with deep learning. Nat Neurosci 21, 1281–1289 (2018).

91. Nath, T., et al. Using DeepLabCut for 3D markerless pose estimation across species and behaviors. Nat Protoc 14, 2152–2176 (2019).

92. Chen, G., et al. Distinct reward processing by subregions of the nucleus accumbens. Cell Rep 42, 112069 (2023).

93. Lafferty, C.K., Yang, A.K., Mendoza, J.A. & Britt, J.P. Nucleus Accumbens Cell Type- and Input-Specific Suppression of Unproductive Reward Seeking. Cell Rep 30, 3729–3742 e3723 (2020).

94. Soares-Cunha, C., et al. Nucleus accumbens medium spiny neurons subtypes signal both reward and aversion. Mol Psychiatry 25, 3241–3255 (2020).

